# Compact red-shifted near-infrared fluorescent proteins enable deep-tissue SWIR imaging with *in vivo* optical clearing

**DOI:** 10.64898/2026.07.25.740731

**Authors:** Kyrylo Yu. Manoilov, Yirui Xu, Jinhuan Luo, Olena S. Oliinyk, Erin Carey, Kateryna O. Tokarchuk, Jinghang Zhang, Guosong Hong, Axel Nimmerjahn, Junjie Yao, Vladislav V. Verkhusha

## Abstract

Compact fluorescent proteins (FPs) with red-shifted emission are needed for deep-tissue short-wavelength infrared (SWIR) imaging. We engineered a GAF domain from the JSC1 cyanobacteriochrome of thermophilic *Leptolyngbya* sp. into three monomeric, biliverdin-binding FPs of 19.1 kDa: miRFP729nano, miRFP732nano and miRFP735nano, with excitation/emission maxima of 714/729, 716/732 and 719/735 nm, respectively. Their off-peak fluorescence beyond 1,000 nm was several-fold higher than that of miRFP718nano previously used for SWIR imaging. miRFP732nano functioned as a fusion tag, a component of target-stabilized nanobodies, and a reporter of NF-κB and AP-1 transcriptional activities. It enabled single-laser, dual-color three-photon imaging with EGFP to depths of ∼950 μm in cortex and ∼300 μm in spinal cord. In mice, miRFP732nano supported SWIR imaging of skeletal muscle, inflammatory signaling and intracellular targets. Combining SWIR detection with biocompatible 4-aminoantipyrine-based *in vivo* tissue clearing enhanced signal and image sharpness. These red-shifted NIR FPs expand the genetically encoded toolkit for deep-tissue imaging.

## Introduction

Optical imaging enables noninvasive visualization of biological structures and processes in living organisms, but its penetration depth is fundamentally limited by light attenuation in tissues. In the visible range (350–650 nm), strong absorption by hemoglobin and melanin, together with extensive light scattering and tissue autofluorescence, restricts both imaging depth and contrast. Shifting excitation and emission to the near-infrared (NIR) range reduces these effects and thereby improves optical imaging at greater depths^1, 2^. In particular, the second NIR window (NIR-II; 1,000–1,700 nm), also termed the short-wavelength infrared (SWIR) range, has attracted substantial interest because it can provide greater penetration depth, improved spatial resolution and higher image contrast than the NIR-I range (650–900 nm)^3–6^. Genetically encoded fluorescent probes with detectable SWIR emission are therefore needed for deep-tissue imaging. Unlike externally delivered fluorophores, NIR fluorescent proteins (FPs) can be expressed in selected cell populations, organelles or protein fusions, used as reporters of biological activity, and multiplexed with visible FPs, biosensors and optogenetic tools^7^.

Most genetically encoded NIR FPs have been engineered from two classes of soluble bacterial photoreceptors: bacteriophytochromes (BphPs) and cyanobacteriochromes (CBCRs). The smallest BphP-based NIR FPs contain at least PAS and GAF domains and have molecular weights of ∼35 kDa ^7^, whereas CBCR-based NIR FPs can consist of a single GAF domain of only 17–19 kDa ^8^. Both classes can utilize the linear tetrapyrrole biliverdin IXα (BV), an intermediate of heme catabolism that is enzymatically produced in mammalian cells by heme oxygenase. BphPs naturally bind BV, whereas most CBCRs bind phycocyanobilin, which is not produced in mammalian cells. Ttherefore, efficient BV binding by CBCR-derived FPs generally requires protein engineering. Nevertheless, the small size and monomeric architecture of CBCR GAF domains have enabled development of compact NIR FPs, including miRFP670nano^9^, miRFP670nano3^10^, miRFP704nano^11^ and miRFP718nano^12^. More recently, a phylogenetically distinct group of CBCRs with substantially red-shifted photochemistry was described, and several of their GAF domains were shown to bind not only phycocyanobilin but also BV^13^. These domains may therefore provide new templates for extending the NIR FP palette toward longer wavelengths. Despite more than one and half decades of NIR FP engineering, the emission maxima of practical NIR FPs have remained at ∼720 nm or below^14, 15^. Consequently, their strongest fluorescence lies within the NIR-I range, and SWIR imaging relies on detection of the relatively weak long-wavelength tails of their emission spectra^12, 16–19^. FP complexes with emission maxima closer to or within the SWIR range have been reported, including the approximately 250 kDa IRFP897 and 350 kDa IRFP1032 complexes, with excitation/emission maxima of 878/897 nm and 1,008/1,032 nm, respectively^20^. However, these multimeric complexes use bacteriochlorophylls *a* or *b*, which are not efficiently produced in mammalian cells, and their large size precludes their general use as protein tags or genetically encoded reporters. Synthetic fluorophores can emit farther into SWIR; for example, CO-1080 exhibits excitation/emission maxima of 1,040/1,054 nm when bound to human serum albumin^21^. Such probes, however, require delivery into tissues, cannot directly report gene expression or transcriptional activity, and do not provide the cell-type and protein-level specificity afforded by genetically encoded FPs. Thus, although the fluorescence maxima of BV-binding NIR FPs remain within NIR-I, their off-peak emission currently provides the most practical genetically encoded approach to SWIR imaging in mammalian systems^12, 22^.

Red-shifting the fluorescent probe addresses only one barrier to deep-tissue imaging, because light-scattering remains substantial even at SWIR wavelengths^23, 24^. Optical tissue clearing provides a complementary strategy by reducing the refractive-index mismatch between aqueous and lipid-rich tissue components, which is a major source of light scattering^25, 26^. Conventional clearing methods generally require high concentrations of harsh chemicals and are therefore restricted largely to fixed or non-living tissues^27, 28^. By contrast, strongly absorbing small molecules, including tartrazine and 4-aminoantipyrine, can increase the refractive index of aqueous tissue components and induce reversible optical transparency in living tissues at biocompatible concentrations^29^. Indocyanine green has also been applied to optical clearing of lipid-rich tissues in the SWIR range, enabling *in vivo* tissue clearing at approximately 1,300 nm^30^. These findings suggest that development of more red-shifted NIR FPs and modulation of the tissue optical environment could be combined to improve deep-tissue fluorescence imaging.

Here, we pursued both approaches. We engineered the BV-binding JSC1_58120g3 GAF domain from a CBCR of thermophilic *Leptolyngbya* sp. strain JSC1^13^ into three small monomeric NIR FPs, termed miRFP729nano, miRFP732nano and miRFP735nano. Their fluorescence maxima extend beyond the previous spectral limit of compact genetically encoded NIR FPs, and their off-peak emission provides substantially greater fluorescence beyond 1,000 nm than miRFP718nano. Focusing on miRFP732nano, we demonstrate its use as a terminal and internal protein tag, a component of target-stabilized NIR fluorescent nanobodies, and a reporter of NF-κB and AP-1 transcriptional activities. We further show single-laser, dual-color multiphoton imaging of miRFP732nano and GFP in the mouse brain and spinal cord. At the macroscopic scale, miRFP732nano enables noninvasive SWIR imaging of skeletal muscle, inflammatory signaling and intracellular targets in living mice. Finally, combining SWIR detection with biocompatible, 4-aminoantipyrine-based *in vivo* tissue clearing further improves fluorescence signal and image sharpness, establishing a complementary strategy for extending genetically encoded imaging deeper into living mammalian tissues.

## Results

### Engineering JSC1_58120g3 into NIR FPs

Directed molecular evolution of the CBCR JSC1_58120g3 GAF domain was performed in BV-producing *E. coli* strain LMG194 co-expressing the *Bradyrhizobium* HO (hmuO) ^18, 31^ and consisted of rational design and random mutagenesis steps (**Supplementary Fig. 1**). The first step of rational design focused on eliminating photoswitching in WT JSC1_58120g3. A single mutation, His117Phe, abolished the photoswitching, but induced a blue shift in the fluorescence emission. To restore the original fluorescence properties, two mutations were introduced: Pro71Cys and Cys116Ser. These substitutions repositioned the cysteine residue responsible for covalent attachment of BV from position 116 to 71. The resulting mutant with 3 substitutions and emission at 732 nm was applied to random mutagenesis.

Following next seven rounds of directed molecular evolution (**Supplementary Fig. 1**), bright in in HeLa cells mutants were obtained that also exhibited blue shift. The Tyr87Phe substitution restored the fluorescence peak at 732 nm but decreased 40% efficient (cellular) brightness in HeLa. Therefore, additional rounds of random mutagenesis followed. Interestingly, the Phe87Tyr mutation reappeared in later rounds of evolution, again producing blue shift and was retained due to increased fluorescence intensity in a miRFP729nano variant.

The step 3 of rational design involved truncating the N-terminus by first 16 residues (removing the N-terminal α-helix) together with the last 5 C-terminal residues to shorten the distance between the C-and N-termini to facilitate future use in internal protein tagging. To stabilize the structure after truncations, nine substitutions were introduced (Phe22Lys, Ala25Leu, Val29Ile, Gly34Leu, Gln85Arg, Glu101Asp, Leu113Ser, Val166Thr, and Ala173Ile). In step 4 of rational design, surface-exposed Cys111 was substituted with Arg to prevent cysteine-mediated crosslinking, which may limit the FP use in oxidative environment^17^. The step 5 of rational design introduced Gln107Arg, Ser111Arg, Cys129Thr, Arg136Glu, Met143Leu, and Val89Thr mutations to maximize emission red-shift of a miRFP735nano variant.

The 16 rounds of directed molecular evolution in total yielded the miRFP732nano variant, with closely located C-and N-termini and containing 25 amino acid amino acid substitutions compared to the WT JSC1_58120g3: Phe22Lys, Thr25Leu, Val29Ile, Phe33Leu, Gly34Asp, Thr53His, Glu56Lys, Gly58Ser, Ser66Ile, Pro71Cys, Gln82Arg, Gln85Arg, Glu101Asp, Phe104Tyr, Arg111Ser, Thr114Ser, Cys116Ser, His117Phe, Glu119Arg, Ser123Arg, Gln136Arg, Leu143Met, Asn149His, Val166Thr, and Ala173Ile (**Supplementary Fig. 1**).

Compared to miRFP732nano, the miRFP729nano variant lacks the Arg111Ser and Gln136Arg substitutions but contains three additional mutations: Tyr87Phe, Leu94Gln, and Val131Ala (26 mutations in total). The miRFP735nano variant lacks mutations Arg111Ser, Gln136Arg, and Leu143Met but has Val89Thr, Gln107Arg, Cys129Thr, and Val131Ala substitutions. All three JSC1_58120g3-derived NIR FPs contain 163 amino acid residues and have calculated molecular weight of 19.1 kDa.

### Characterization of the developed NIR FPs *in vitro*

In BV-producing bacteria brightness of miRFP729nano and miRFP732nano was comparable while brightness of miRFP735nano was notably lower (**Supplementary Fig. 2**). We next demonstrated that red-shifted properties of miRFP732nano enable its spectral separation with previously reported other CBCR-derived NIR FPs, such as miRFP670nano3 ^10^, miRFP704nano ^11^, and miRFP718nano ^12^, using standard spectral unmixing approach (**Supplementary Fig. 3**), thus allowing spectral multiplexing.

The absorption spectra of the developed red-shifted NIR FPs exhibited minor peaks at 384 nm, corresponding to the Soret band, characteristic of tetrapyrrole-binding proteins. Major Q-band peaks were observed at 714, 716, and 719 nm for miRFP729nano, miRFP732nano, and miRFP735nano, respectively (**Fig. 1a**). The fluorescence excitation maxima were similar to those of the absorbance Q-bands (**Fig. 1b**). The corresponding emission maxima were 729, 732, and 735 nm (**Fig. 1c**, and **Supplementary Table 1**). The molar extinction coefficients of miRFP729nano, miRFP732nano, and miRFP735nano were 69,300 M⁻¹ cm⁻¹, 64,400 M⁻¹ cm⁻¹, and 59,200 M⁻¹ cm⁻¹, and fluorescence quantum yields were 1.9%, 1.6%, and 1.0%, respectively **Supplementary Table 1**).

**Figure 1.**
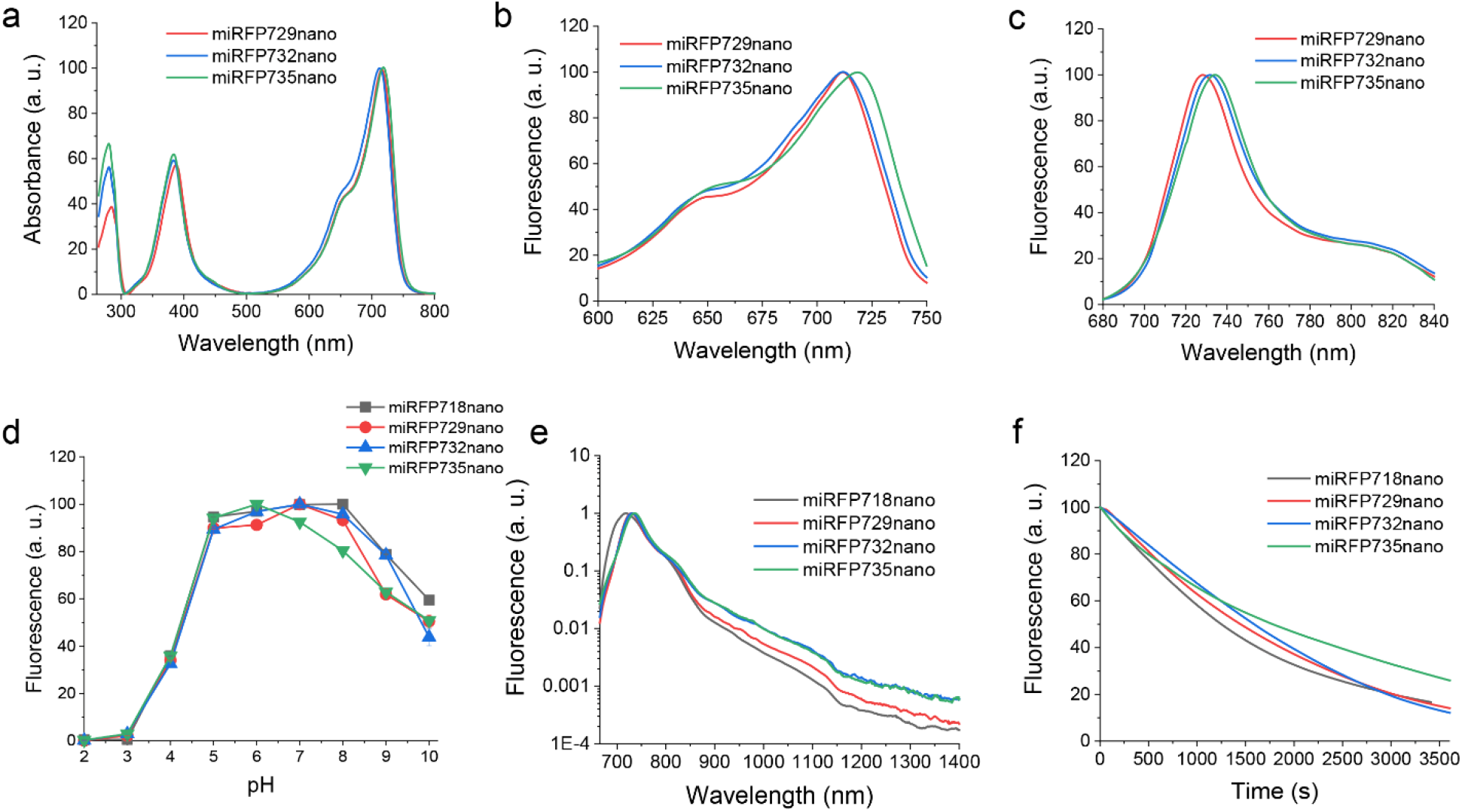
Properties of the engineered miRFP729nano, miRFP732nano and miRFP735nano. (**a**) Absorbance spectra. (**b**) Fluorescence excitation spectra. (**c**) Fluorescence emission in visible spectral range. (**d**) pH dependencies of fluorescence. (**e**) Fluorescence emission spectra extended to 1400 nm. (**f**) Photobleaching kinetics in live HeLa cells. In (d)-(f), the most red-shifted CBCR-derived single-domain NIR FP, miRFP718nano, was used for benchmarking.

The pH-dependence of fluorescence for three engineered proteins was consistent with that of previously described CBCR NIR FPs (**Fig. 1d**), exhibiting apparent p*K*a values of 4.2 for miRFP729nano and miRFP732nano, and 4.1 for miRFP735nano (**Supplementary Table 1**). Protein gel electrophoresis indicates that miRFP732nano behaves as a monomer (**Supplementary Fig. 4**).

To estimate potential performance of the new NIR FPs in SWIR imaging, we compared their fluorescence emission beyond 1000 nm to that of miRFP718nano (**Fig. 1e**), which has been previously shown to perform the best among NIR FPs in a “off-peak emission” SWIR imaging.^12^ All three engineered proteins exhibited substantially higher relative fluorescence in this range. Relative to miRFP718nano, the fluorescence intensity above 1000 nm was 1.9-fold higher for miRFP729nano, and over 3-fold higher for miRFP732nano and miRFP735nano (**Supplementary Table 1**).

To further test whether this enhanced SWIR emission was translated into improved imaging, we compared purified miRFP732nano and miRFP718nano at equal concentrations in a silicone tube tissue phantom, imaged under both NIR-I and SWIR detection (**Supplementary Fig. 5**). Under NIR-I detection, miRFP718nano yielded ∼1.5-fold stronger fluorescence than miRFP732nano, consistent with its higher intrinsic brightness. This relationship reversed under SWIR detection, where miRFP732nano outperformed miRFP718nano by ∼2.1-fold. This advantage in absolute signal is smaller than the 3-fold difference in peak-normalized SWIR emission (**Fig. 1e**), as expected: the phantom measures total detected signal at equal protein concentration, so the higher intrinsic brightness of miRFP718nano partially offsets the larger SWIR-tail fraction of miRFP732nano, and the fixed 680 nm excitation preferentially excites miRFP718nano (excitation maximum 690 nm) over miRFP732nano (716 nm). Thus, despite its lower overall brightness, the red-shifted emission of miRFP732nano was the stronger candidate for the SWIR range, motivating its use *in vivo*.

### Performance of miRFP732nano in live mammalian cells

All three new NIR FPs exhibited good effective brightness in mammalian cells without supply of exogeneous BV. 72 h after transfection, miRFP732nano cells were ∼3.9 times less bright than miRFP718nano. miRFP729nano was 36% brighter than miRFP732nano and miFRP735nano was 71% less bright than miRFP732nano (**Supplementary Table 1**). At the same time, the proteins exhibited the higher photostability in live mammalian cells relative to other NIR FPs, with photobleaching half-times of 1445, 1585, and 1775 s (**Fig. 1f** and **Supplementary Table 1**).

To evaluate the performance of miRFP732nano as a protein tag, we constructed a set of fusion proteins with miRFP732nano tag either fused terminally or internally (**Fig. 2a**). In live HeLa cells, C-terminal fusion with β-actin displayed proper filamentous localization. N-terminal fusions with a mitochondria-targeting sequence and with histone H2B resulted in expected localization to mitochondria and chromatin, respectively. To assess the performance of miRFP732nano as an internal tag, we inserted it between the helical and GTPase domains of the G-protein α-subunit (Gαs) and into the intracellular loop 3 of the β2-adrenergic receptor (β2AR). In both cases, internal miRFP732nano fusions exhibited correct membrane localization, consistent with proper folding and function of the fusion proteins. These results demonstrate that miRFP732nano performs well as the terminal and internal tag, supporting its applicability to label diverse cellular proteins.

**Figure 2.**
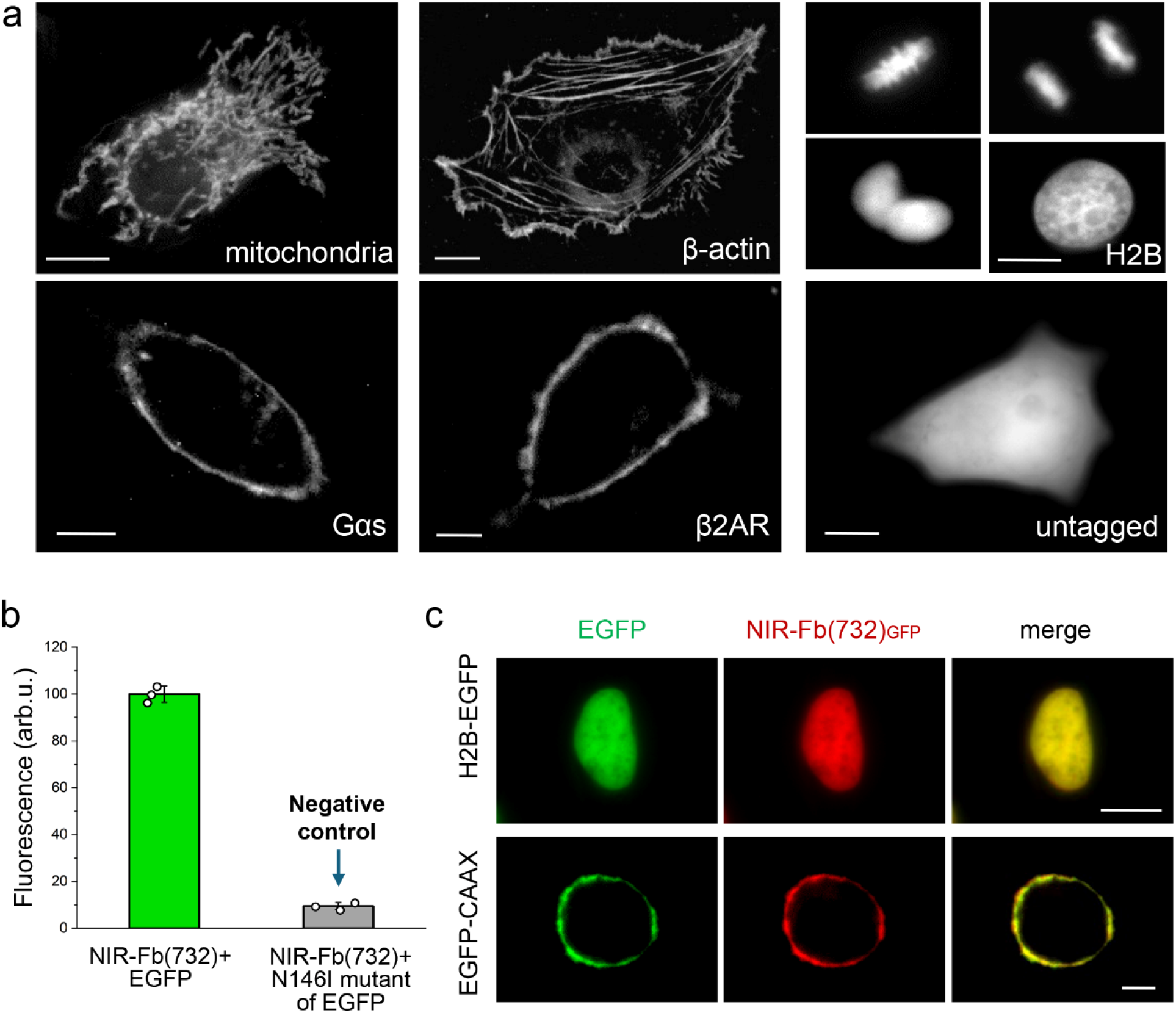
Performance of miRFP732nano in live HeLa cells. (**a**) C-terminal fusion: β-actin; N-terminal fusions: mitochondrial signal and histone H2B; internal fusions: β2 adrenergic receptor (β2AR) and G-protein α subunit (Gαs). Ex. 685/20 nm, and em. 725/40 nm. Scale bars, 10 μm. (**b-c**) miRFP732nano was inserted into the anti-GFP nanobody, resulting in the NIR-Fb(732) target-stabilizable nanobody construct. (b) Flow cytometry. When a specific intracellular target, such as EGFP, is present, NIR-Fb(732) is stabilized by binding to the target, preserving miRFP732nano fluorescence. When there is no specific target (Negative control: EGFP/N146I mutant is not recognized by anti-GFP nanobody), NIR-Fb(732) is destabilized and fast degraded. EGFP was excited with 488 nm and detected with 525/30 nm, miRFP732nano was excited with 640 nm and detected with 725/40 nm. (c) Epifluorescence microscopy. NIR-Fb(732) selectively targets EGFP-fused histone H2B (top) and membrane-anchored EGFP-CAAX (bottom), resulting in NIR fluorescence colocalizing with EGFP signal. EGFP was imaged using ex. 485/20 nm and em. 525/30 nm, and miRFP732nano was imaged using ex. 685/20 nm and em. 725/40 nm. Scale bars, 10 μm.

### Development of miRFP732nano-based NIR-Fb

Recently a target-stabilizable NIR fluorescent nanobody (Fb) approach has been developed ^10, 32^. In this technique, a CBCR-derived NIR FP, such as miRFP670nano and miRFP718nano, with closely located termini is inserted at a specific position into a nanobody scaffold. When expressed in mammalian cells, the resulting NIR-Fb is stabilized by binding of a cognate intracellular target and is rapidly degraded in the absence of the target, thus producing no fluorescence in target-negative cells. To test whether miRFP732nano is compatible with the NIR-Fb architecture, we inserted miRFP732nano into an anti-GFP nanobody, resulting in NIR-Fb(732)_GFP_ of ∼35 kDa construct, and evaluated it in live cells (**Fig. 2b,c**). In the absence of the EGFP target, NIR-Fb(732)_GFP_ fluorescence was not detected, consistent with proteasomal degradation of the unbound form ^10, 32^. In contrast, co-expression of EGFP resulted in strong NIR fluorescence, indicating target-dependent stabilization of the construct. Furthermore, NIR-Fb(732)_GFP_ selectively labeled EGFP-tagged proteins in various compartments, including nuclear histone H2B and membrane-anchored EGFP-CAAX (**Fig. 2c**). This demonstrated the miRFP732nano compatibility with the NIR-Fb platform that could be used to develop other NIR-Fbs functional in live cells.

### Development of the inflammation reporters

To evaluate miRFP732nano in monitoring cell signaling, we engineered miRFP732nano-based transcription reporters for two key inflammation signaling pathways involving nuclear factor-κB (NF-κB) and activator protein 1 (AP-1) (**Fig 3**). After activation, NF-κB and AP-1 bind to their respective consensus response DNA elements, initiating transcription of downstream target genes. To assess the performance of the engineered miRFP732nano-based transcription reporters, we transiently transfected HEK293T cells. Non-stimulated cells exhibited weak or no miRFP732nano fluorescence, indicating low basal activity. Upon stimulation with tumor necrosis factor-α (TNF-α), cells transfected with the NF-κB reporter displayed strong NIR fluorescence (**Fig. 3a–c**). Dose-response analysis showed that 5 ng/mL TNF-α fully activated the reporter, with EC_50_ of ∼1 ng/mL (**Fig. 3c**). Stimulation of cells transfected with the AP-1 reporter with PMA induced robust miRFP732nano expression at 50 ng/mL, with EC_50_ of ∼5 ng/ml (**Fig. 3d-f**).

**Figure 3.**
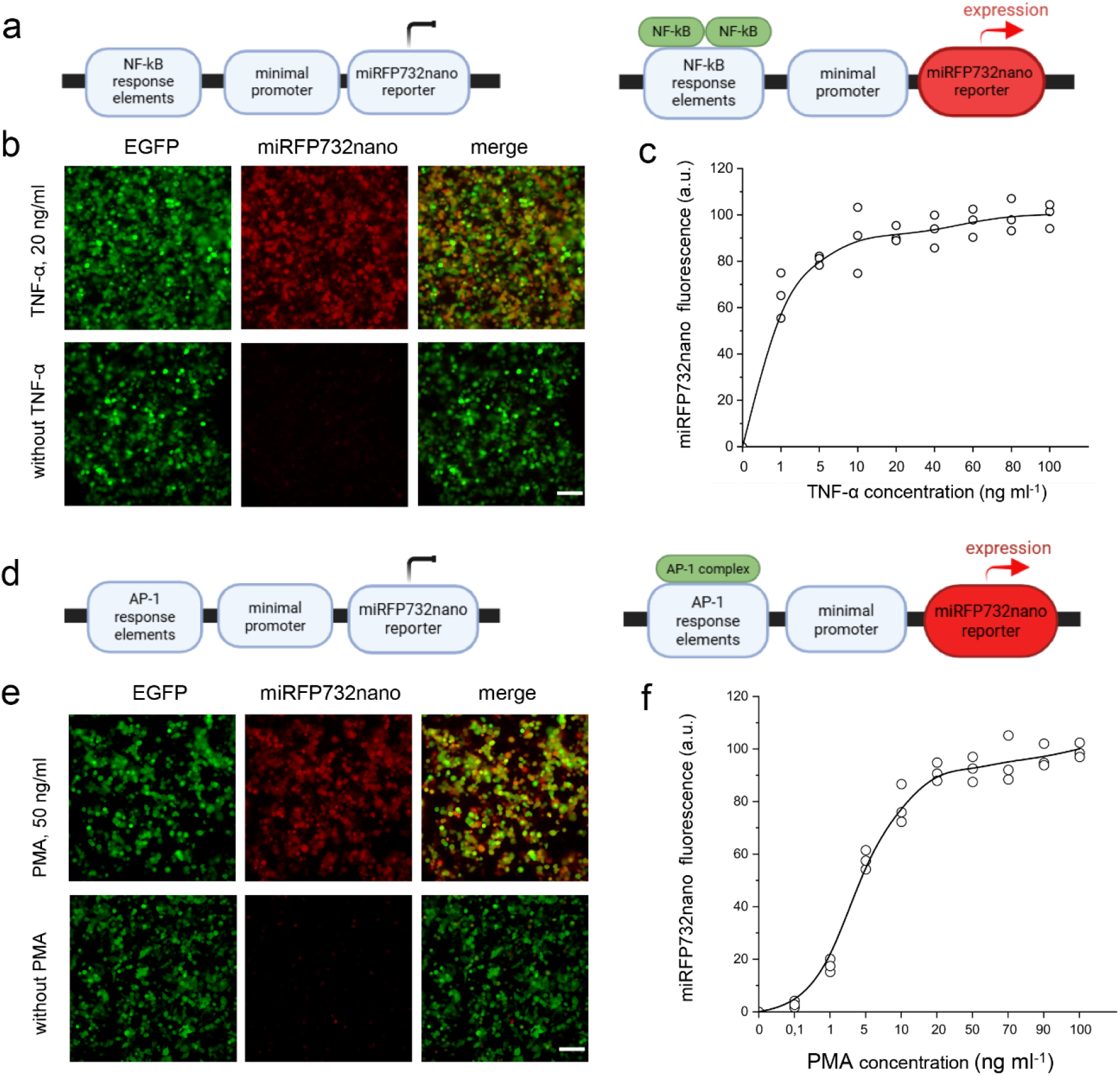
Performance of miRFP732nano-based NF-kB and AP-1 inflammation reporters in cells. **(a)** Schematic of the miRFP732nano-based NF-κB transcription reporter. **(b)** Fluorescence images of live HEK293T cells transfected with the NF-κB reporter and co-transfected with pEGFP-N1 (1:10). Cells were either stimulated with TNF-α (20 ng mL⁻¹, top row) or left untreated (bottom row). **(c)** TNF-α dose–response of the NF-κB reporter measured by flow cytometry of live HEK295T cells. Data are presented as mean ± s.d. for *n* = 3 independent transfections. **(d)** Schematic of the miRFP732nano-based AP-1 transcription reporter. **(e)** Fluorescence images of live HEK293T cells transfected with the AP-1 reporter and co-transfected with pEGFP-N1 (1:10). Cells were either stimulated with PMA (50 ng mL⁻¹, top row) or left untreated (bottom row). **(f)** PMA dose–response of the AP-1 reporter measured by flow cytometry using live HEK293T cells. Data are presented as mean ± s.d. for *n* = 3 independent transfections. In (b) and (e), epifluorescence microscopy: EGFP and miRFP732nano were imaged using 485/20 nm excitation and 525/30 nm emission filters, and 685/20 nm excitation and 725/40 nm emission filters, respectively. Scale bars, 100 µm. In (c) and (f), flow cytometry: EGFP was excited with 488 nm and detected with 525/30 nm, and miRFP732nano was excited with 640 nm and detected with 725/40 nm filter. Representative images from three independent experiments are shown.

### Multiphoton imaging in the brain and spinal cord *in vivo*

To evaluate miRFP732nano performance in neurons in living mice, we subcloned it into an AAV vector carrying a hSyn promoter and the soma-targeting peptide RiboL1.^33, 34^ When fused with RiboL1 that binds ribosomes, miRFP732nano is translated by ribosomes already active in the cytoplasm, thus efficiently escaping the nuclei (**Supplementary Fig. 6**). To increase BV level in neurons *in vivo*, we followed the approach in which inhibition of BV reductase A by shRNA was combined with co-expression of HO1 (human *HMOX1*), ferredoxin (Fd), and ferredoxin–NADP⁺ reductase (Fnr) ^35,36^ The shRNA was expressed from the H1 promoter while HO1, Fd, and Fnr were produced as a single transcript under a short CAG promoter. This strategy increased the miRFP732nano fluorescence in mouse NIH3T3 cells by ∼2.2-times 48 h post-transfection, which was only ∼30% smaller than that obtained by supplying cells with 25 μM exogenous BV (**Supplementary Fig. 7**). The size of the DNA construct including both ITR regions was 4577 bp, allowing incorporation into AAV.

We co-injected both AAV vectors (1:1 mix; **Methods**) into the somatosensory cortex or the spinal dorsal horn of *Cx3cr1^GFP/+^*mice, which express GFP in microglia, the central nervous system’s resident immune cells, allowing assessment of dual-color imaging capabilities and innate immune responses, as in our previous work.^10^ One to two months after injection, animals were prepared for *in vivo* imaging to determine protein expression and achievable imaging depth. We previously demonstrated^10^ dual-color two-photon (2P) imaging of GFP and the less red-shifted miRFP670nano3 (ex./em. 645/670 nm) using a single two-photon laser to depths of ∼870 μm and ∼120 μm from the cortical and spinal surfaces, respectively. miRFP732nano extends the *in vivo* imaging range further into NIR. However, the increased spectral separation for excitation of GFP and miRFP732nano presents challenges for multicolor 2P imaging, and near-surface background generation can degrade the signal-to-background ratio^37^ at greater 2P imaging depths. Multicolor three-photon (3P) microscopy is highly desirable for expanding deep-tissue imaging applications, but it relies on suitable, red-shifted FPs. We found that 3P microscopy permitted the simultaneous visualization of miRFP732nano-expressing neurons and GFP-positive microglia in *Cx3cr1^GFP/+^*mice at depths up to ∼950 μm in the cortex and up to ∼300 μm in the spinal dorsal horn using 1300 nm excitation with a single laser (**Fig. 4**, and **Methods**). As expected from using RiboL1, miRFP732nano was mainly confined to neuronal cell bodies (**Fig. 4**, and **Supplementary Fig. 8**). Surrounding microglia showed no overt signs of morphological activation, indicating that miRFP732nano was well-tolerated under the expression conditions used in this study. These data suggest that miRFP732nano has sufficient 3P cross-section to enable single-laser dual-color 3P *in vivo* imaging.

**Figure 4.**
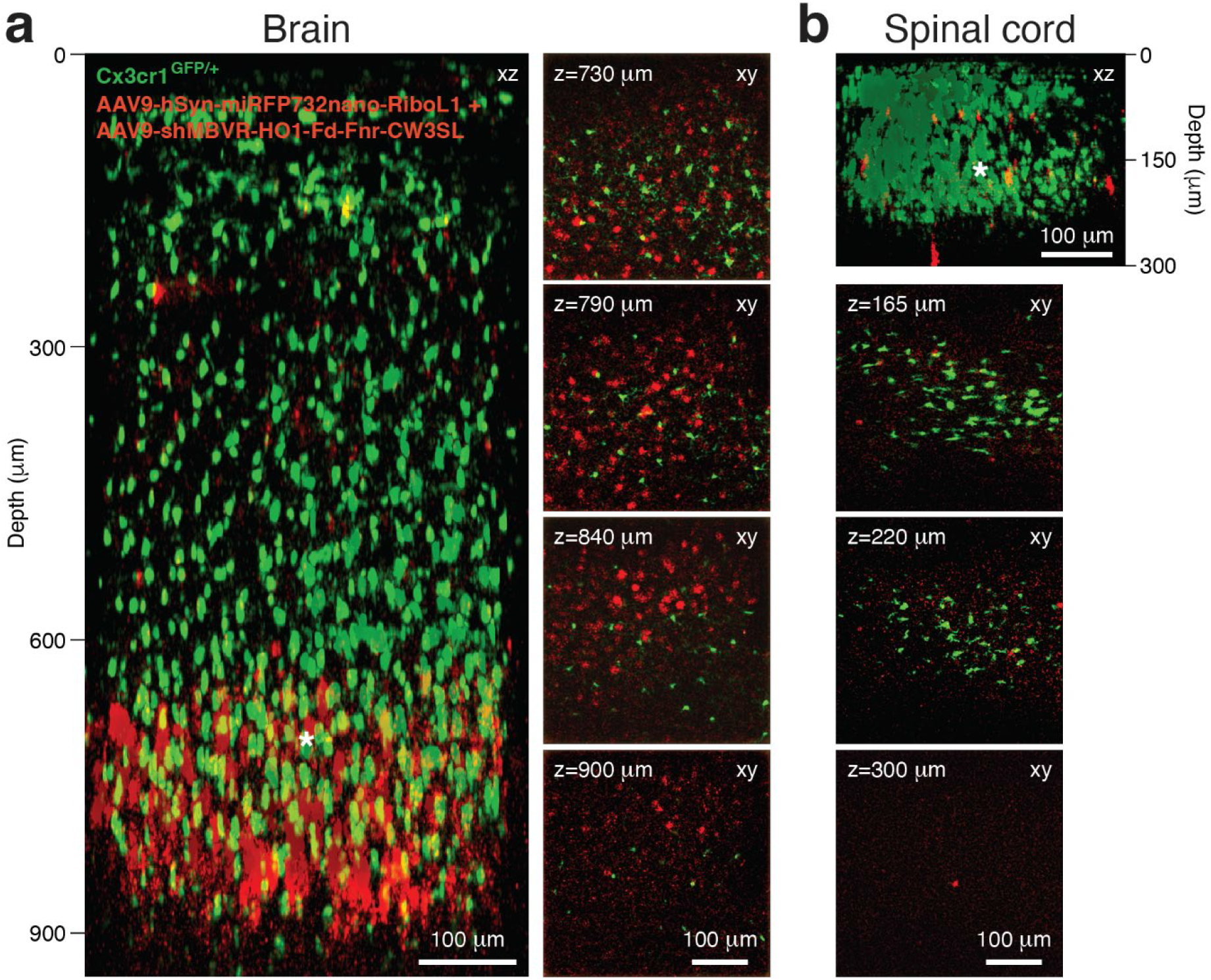
Multiplex three-photon imaging with miRFP732nano in the brain and spinal cord of live *Cx3cr1^GFP/+^* mice. (**a**) *Left*, xz projection from an xy fluorescence image stack showing neuronal somata (red) and microglia (green) in the somatosensory cortex of a 17.5-week-old *Cx3cr1^GFP/+^*mouse, 4.5 weeks after stereotactic co-injection of AAV9-hSyn-miRFP732nano-RiboL1 and AAV9-shMBVR-HO1-Fd-Fnr-CW3SL viruses into deep cortical layers. *Right*, four example images from the xy fluorescence image stack at the indicated depths (z) from the pial surface. Scale bars, 100 μm. (**b**) *Top*, xz projection from an xy fluorescence image stack showing neuronal somata (red) and microglia (green) in the lumbar spinal cord of an 8.5-week-old *Cx3cr1^GFP/+^*mouse, 4 weeks after stereotactic co-injection of AAV9-hSyn-miRFP732nano-RiboL1 and AAV9-shMBVR-HO1-Fd-Fnr-CW3SL viruses into superficial dorsal horn laminae. *Bottom*, three example images from the xy fluorescence image stack at the indicated depths (z) from the pial surface. GFP and miRFP732nano fluorescence images were acquired simultaneously with 1300 nm excitation. Scale bars, 100 μm. White asterisks mark the AAV injection depth. Representative images from three brain and four spinal cord imaging experiments are shown.

### SWIR imaging through scattering media with tissue clearing

Although 3P and SWIR imaging operate at longer wavelengths, light-scattering remains a major barrier to deep-tissue imaging. We, therefore, further explored combination of SWIR detection with optical clearing as a complementary strategy to suppress tissue scattering. We first evaluated several reported optical clearing agents for SWIR imaging, including ICG ^30^, tartrazine ^29^, and 4-aminoantipyrine ^29, 38^ (**Supplementary Fig. 9**). ICG has strong absorption within the NIR-I region (∼700-850 nm), making it incompatible with NIR-I imaging. While both tartrazine and 4-aminoantipyrine produced high optical transparency within the NIR-I and SWIR spectral regions, tartrazine presented more prominent precipitation at high concentrations, compromising its practical utility (**Supplementary Fig. 9c,d**). In contrast, 4-aminoantipyrine maintained high optical transparency and physical stability and was therefore selected for further *in vivo* studies.

To evaluate the optical clearing effect in SWIR region, we used a SWIR-emitting BIBDAH dye ^39^ (**Supplementary Fig. 9a**) to generate a letter-patterned phantom and covered it with a scattering layer consisting of intralipid and water. Under these conditions, the letter pattern was undiscernible due to strong optical scattering. Upon dying the scattering layer with 4-aminoantipyrine (380 mg/ml) while keeping the layer thickness unchanged, optical transparency was increased, enabling clear visualization of the letter pattern (**Fig. 5a, b**). These results confirm that 4-aminoantipyrine effectively reduces optical scattering in the SWIR region.

**Figure 5.**
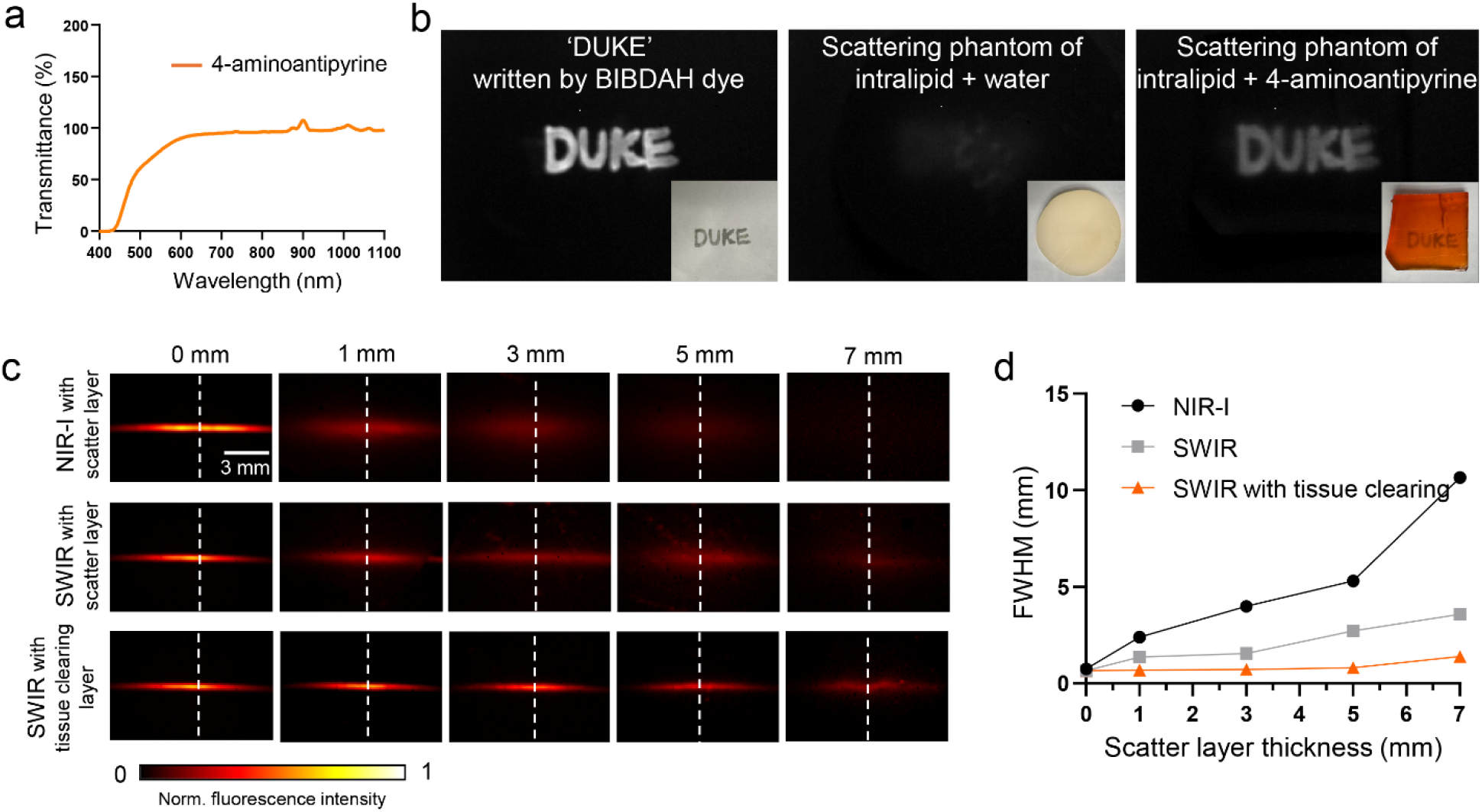
Combined detection in SWIR region and 4-aminoantipyrine-based tissue clearing improve fluorescence imaging in scattering media. **(a)** Measured optical transmittance spectrum of 4-aminoantipyrine, showing high optical transmittance across the NIR-I and SWIR spectral regions. **(b)** Demonstration of the optical clearing effect in the SWIR region. BIBDAH dye was used to write ‘DUKE’ on a substrate and imaged under SWIR illumination (1050 nm excitation, 1100 nm long-pass emission). The pattern was obscured by an intralipid + water scattering layer (middle) but became clearly visible when the water was replaced with solution of 4-aminoantipyrine (right). Insets show white-light photographs of the corresponding samples. **(c)** NIR-I and SWIR fluorescence imaging of the silicone-tube phantom filled with 5 mg/mL of the purified miRFP732nano protein, covered by scattering media of increasing thickness (0–7 mm). *Top row:* NIR-I imaging through scattering layers. *Middle row:* SWIR imaging through scattering layers. *Bottom row:* SWIR imaging through tissue-clearing layers containing 4-aminoantipyrine. Scale bar, 3 mm. **(d)** Imaging sharpness as the function of scattering layer thickness for NIR-I, SWIR, and SWIR with tissue clearing conditions, quantified as the full width at half maximum (FWHM) of the tube intensity profile along the white dashed lines in (c).

We next studied whether tissue clearing could enhance fluorescence imaging of miRFP732nano through scattering media. Purified miRFP732nano was loaded into a silicone tube phantom and imaged under three conditions: (i) NIR-I with a scattering layer, (ii) SWIR with a scattering layer, and (iii) SWIR with a scattering layer after clearing, at increasing layer thicknesses from 0 to 7 mm (**Fig. 5c** and **Supplementary Fig. 10a**). The scattering layer consisted of intralipid and water in condition (i) and (ii) and was cleared by 4-aminoantipyrin as described above in condition (iii). Under conditions (i) and (ii), the NIR-I and SWIR image quality deteriorated progressively with increasing layer thickness. This image degradation was more pronounced in NIR-I where tube profiles became substantially broadened and the boundary was largely lost at 7 mm depth, whereas SWIR imaging preserved sharper tube boundaries throughout the tested depth range. Under condition (iii), 4-aminoantipyrine markedly improved the optical transparency of the scattering layer, yielding the sharpest tube profiles at increasing imaging depths.

To quantitatively evaluate imaging quality, we measured the full width at half maximum (FWHM) of the fluorescence intensity profiles across the tube (**Fig. 5d** and **Supplementary Fig. 10b**). In the absence of a scattering layer, all three conditions yielded comparable FWHM of ∼0.7 mm. As layer thickness increased, FWHM under NIR-I imaging rose sharply, reaching 10.7 mm at 7 mm depth, whereas SWIR imaging provided a FWHM of 3.6 mm at the same depth. Notably, under the tissue clearing condition, the FWHM increased to 1.4 mm at 7 mm depth, representing a 7.7-fold improvement over the NIR-I and 2.6-fold improvement over the SWIR imaging without clearing. These results demonstrate that both SWIR detection and tissue clearing contribute to the reduced optical scattering, with their combination providing the greatest spatial resolution enhancement at depth.

### *In vivo* imaging of miRFP732nano-expressing muscles

To evaluate whether the imaging improvements could be translated to living animals, we performed imaging of mouse hindlimb muscles expressing miRFP732nano after intramuscular transduction of an AAV vector encoding miRFP732nano under short CAG promoter (**Supplementary Fig. 11**). Images were acquired in both the NIR-I and SWIR ranges before and after tissue clearing. The final reference images were captured following skin removal (**Fig. 6a**).

**Figure 6.**
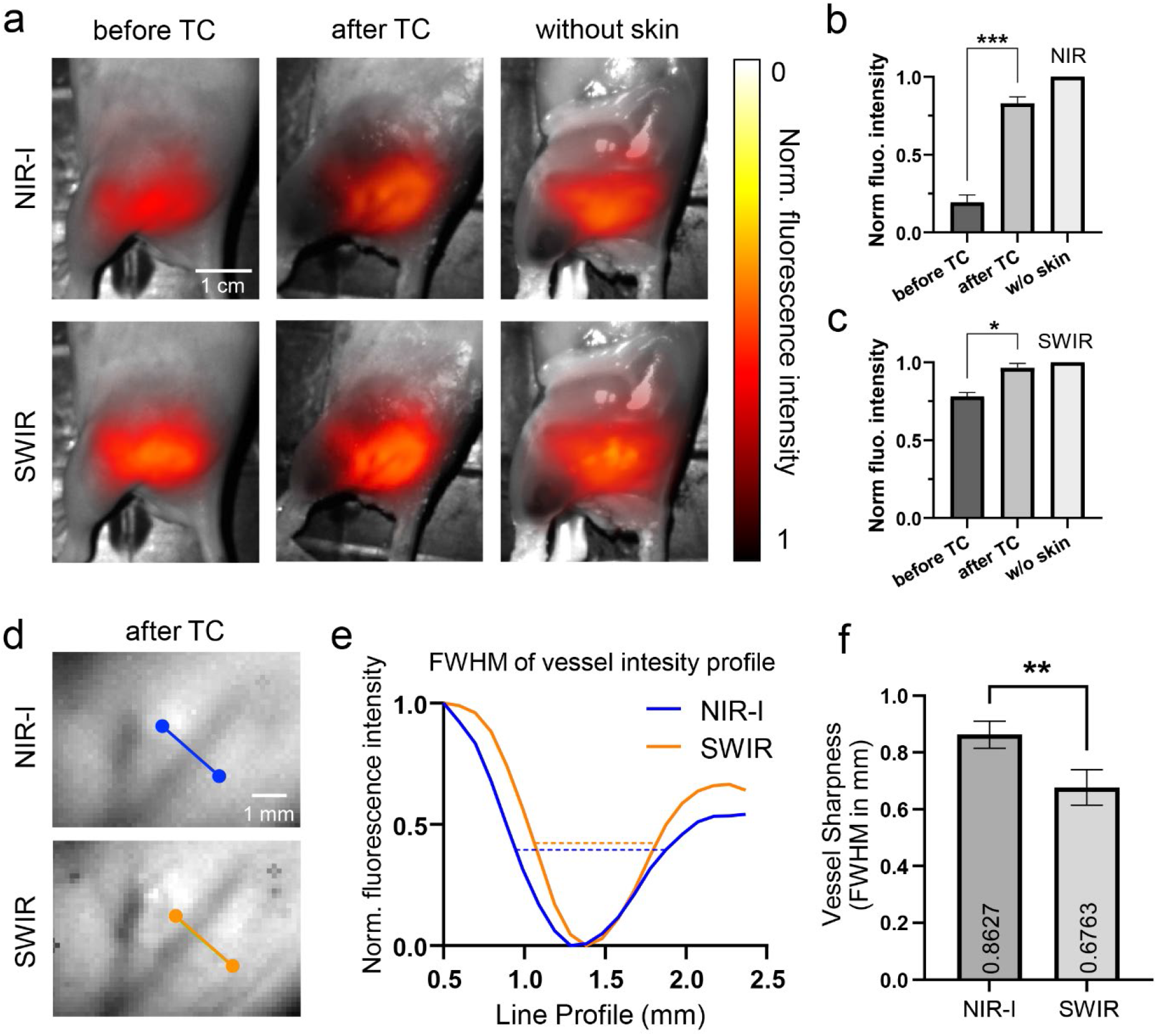
Enhanced imaging of miRFP732nano in hindlimb muscle in SWIR spectral region and with tissue clearing *in vivo*. miRFP732nano was transduced by *in situ* injection of the AAV vector encoding miRFP732nano under CAG promoter. **(a)** Representative fluorescence images of the hindlimb muscle, acquired in the NIR-I and SWIR spectral ranges before tissue clearing (before TC), after tissue clearing (after TC), and after skin removal (without skin). Scale bar, 1 cm. **(b-c)** Quantification of normalized fluorescence intensity in NIR-I and SWIR under three conditions described in (a). Data is presented as mean values ± s.d. (*n* = 3 animals). Statistical significance was assessed using paired two-tailed parametric t-tests: *P < 0.05, **P < 0.01, ***P < 0.001. **(d)** Zoomed-in views of vascular regions after tissue clearing. A representative vessel was selected at matched locations in the NIR-I and SWIR images for direct comparison. Scale bar, 1 mm. **(e)** Fluorescence intensity profiles along the lines indicated in (d), under NIR-I (blue) and SWIR (orange) imaging, quantified by the full width at half-maximum (FWHM); a smaller FWHM indicates a sharper vessel boundary. **(f)** Quantitative comparison of the vessel sharpness in the NIR-I and SWIR spectral ranges based on FWHM. SWIR imaging shows the reduced FWHM relative to NIR-I, indicating sharper vessel delineation (n = 3 vessels, paired t-test, **P < 0.01).

Overall, in both NIR-I and SWIR ranges, tissue clearing enhanced visualization of the underlying muscle tissue, resulting in more distinct structural features. Quantitative analysis confirmed increased fluorescence intensity from the miRFP732nano-expressing muscles following tissue clearing in both NIR-I and SWIR (**Fig. 6b, c**). The relative improvement was greater in the NIR-I window than in the SWIR window, reflecting their different sensitivity to scattering. Compared with NIR-I imaging, SWIR imaging is inherently less affected by tissue scattering, making the benefit of tissue clearing correspondingly smaller. Following tissue clearing, fluorescence signals in both spectral ranges were comparable to those obtained after skin removal, indicating that optical clearing effectively reduced optical scattering in the skin.

To assess image sharpness, we further examined the vascular structures within the cleared tissue. Representative vessels were selected from the NIR-I and SWIR images, and fluorescence intensity profiles were extracted across the vessel lumen (**Fig. 6d, e**). SWIR imaging produced sharper delineation of vascular boundaries than NIR-I. Consistent with this observation, quantitative analysis revealed ∼21% smaller FWHM in SWIR than in NIR-I images (0.68 mm *vs.* 0.86 mm; **Fig. 6f**). Together, the combination of tissue clearing and SWIR enables improved visualization of deep fluorescent signals from the muscles expressing miRFP732nano *in vivo*.

### Longitudinal imaging of miRFP732nano-based transcriptional reporters of inflammation

To evaluate the miRFP732nano-based reporters for monitoring biological processes *in vivo*, we performed longitudinal fluorescence imaging of miRFP732nano reporters of NF-κB and AP-1 expression in the liver, following lipopolysaccharide (LPS)-induced inflammation. The reporter plasmids were delivered to the liver by hydrodynamic transfection, and fluorescence signals were monitored in both the NIR-I and SWIR ranges (**Supplementary Fig. 12**). In both reporter mouse groups, fluorescence progressively increased after the *i.v.* LPS injection, reflecting activation of inflammatory signaling in the liver. Fluorescence signals became detectable in both reporter groups within 6 h after LPS treatment. The NF-κB reporter then increased rapidly and approached a plateau by 24 h (**Fig. 7a, b**), whereas the AP-1 reporter showed a slower, more gradual increase that continued throughout the observation period of 24 h without reaching a clear plateau (**Fig. 7c, d**), consistent with the distinct activation kinetics of the two inflammatory pathways^40^. These data demonstrate that miRFP732nano-based transcriptional reporters enable noninvasive monitoring of the distinct activation dynamics of signaling pathways *in vivo*.

**Figure 7.**
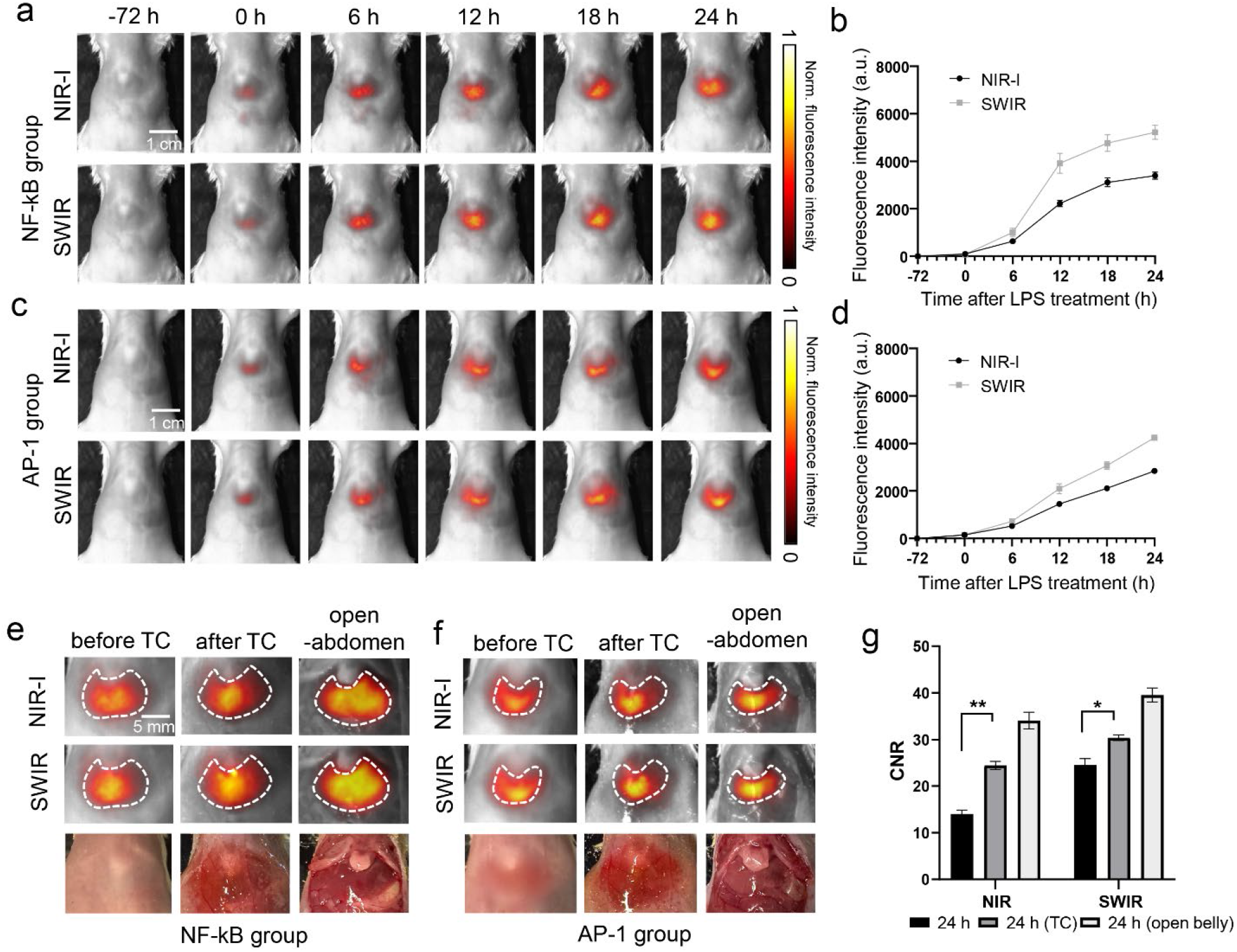
Longitudinal imaging of miRFP732nano-based inflammatory reporters in the mouse liver and effect of tissue clearing *in vivo*. (a, c) Representative NIR-I and SWIR fluorescence images of the NF-κB (a) and AP-1 (c) transcription reporters in the liver at the indicated time points after lipopolysaccharide (LPS) treatment. Scale bar, 1cm. **(b, d)** Fluorescence intensity over time for the NF-κB (b) and AP-1 (d) reporters in the NIR-I and SWIR spectral ranges. **(c)** Zoomed-in fluorescence images of the liver region in NIR-I and SWIR ranges with corresponding regions of interest (ROIs) outlined. Images are shown before and after the tissue clearing, and with mouse belly open. Tissue clearing improves signal visibility in both spectral ranges, with a more pronounced enhancement observed in the NIR-I range (*n* = 3 animals for each group). **(e, f)** Zoomed-in liver images for the NF-κB (e) and AP-1 (f) reporter groups in the NIR-I and SWIR spectral ranges, shown before tissue clearing, after tissue clearing, and after abdominal opening. Regions of interest (ROIs) used for quantification are outlined; bottom rows show corresponding white-light photographs. **(g)** Contrast-to-noise ratio (CNR) quantified from the ROIs in (e, f). Data is presented as mean values ± s.d. (*n* = 3 animals for each group). Statistical significance was assessed using paired two-tailed parametric t-tests: *P < 0.05, **P < 0.01, ***P < 0.001.

To further evaluate the contribution of tissue clearing, fluorescence signals were examined before tissue clearing, after tissue clearing, and after abdominal opening at 24 h after LPS administration (**Fig. 7e,f**). Tissue clearing markedly improved visualization of the liver and increased the contrast-to-noise ratio (CNR) under both NIR-I and SWIR imaging (**Fig. 7g**). The improvement was more pronounced under NIR-I (∼1.7-fold, from CNR 14.0 to 24.4), consistent with the stronger attenuation of NIR-I light by the overlying skin. Notably, SWIR imaging without tissue clearing already achieved a CNR nearly identical to that of NIR-I combined with tissue clearing (24.6 *vs.* 24.4), indicating that the penetration advantage of the SWIR range.

Tissue clearing recovered only a portion of the signals otherwise lost to the abdominal wall. The residual difference is consistent with the diffusion-limited penetration of topically applied clearing agents, which reduce scattering in superficial tissues more effectively than in deeper layers ^29^. Nevertheless, the combination of SWIR detection and tissue clearing offers a practical means to enhance reporter detection in deep tissues.

### Target-dependent activation of miRFP732nano-based NIR-Fb *in vivo*

To determine whether miRFP732nano-based NIR-Fb nanobody retains its target-dependent stabilization *in vivo*, we used EGFP expressed in the mouse liver and delivered the AAV-encoded NIR-Fb(732)_GFP_ systemically via *i.v.* injection (**Fig. 8a**). Fluorescence signals were detected in the liver in both the NIR-I and SWIR windows (**Fig. 8b**). Although the NIR-Fb(732)_GFP_ nanobody was delivered systemically, fluorescence was largely confined to the liver where GFP was expressed. The NIR-I and SWIR images differed markedly in the background autofluorescence: under NIR-I imaging, the liver signal was relatively low, accompanied by strong autofluorescence from the surrounding abdominal tissues and gastrointestinal contents, whereas under SWIR imaging this background was almost absent outside the liver. The same advantage was confirmed after abdominal opening (**Fig. 8c**).

**Figure 8.**
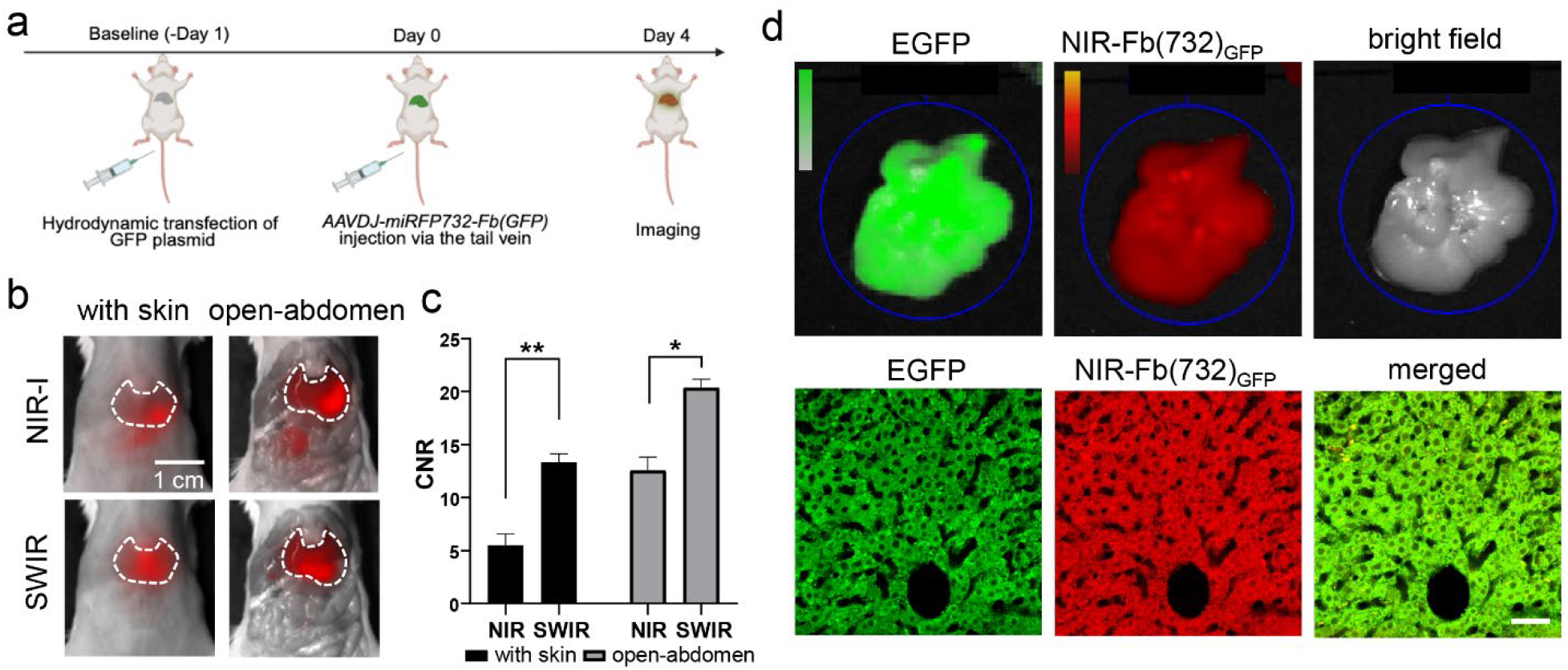
Target-dependent stabilization of miRFP732nano-based NIR-Fb(732)_GFP_ anti-GFP nanobody in the mouse liver *in vivo*. **(a)** Schematic of the experimental workflow. First, EGFP was expressed in the liver by hydrodynamic transfection of the pcDNA-EGFP plasmid. Day later, the EGFP-targeting NIR-Fb(732)_GFP_ nanobody encoded by AAV vector under ubiquitous CAG promoter was *i.v.* administered, and fluorescence imaging was performed 4 days after the AAV transduction. **(b)** Representative NIR-I and SWIR fluorescence images of the liver with intact skin and after abdominal opening. Dashed contours indicate the liver region of interest (ROI). Scale bar, 1 cm. **(c)** Contrast-to-noise ratio (CNR) of liver fluorescence in the NIR-I and SWIR spectral ranges under intact-skin and open-abdomen conditions. Data is presented as mean values ± s.d. (*n* = 3 animals). Statistical significance was assessed using paired two-tailed parametric t-tests: *P < 0.05, **P < 0.01, ***P < 0.001. **(d)** *Ex vivo* evaluation of the NIR-Fb(732)_GFP_ stabilization in EGFP-expressing liver tissue. *Top row:* IVIS Spectrum whole-organ images showing EGFP fluorescence (green; 7.0 ×10⁹ p/s/cm²/sr), miRFP732nano fluorescence (red; 3.4 ×10⁹ p/s/cm²/sr), and the corresponding white-field image. *Bottom row:* confocal images of EGFP (green), miRFP732nano (red), in the merged image. Scale bar, 50 μm.

To further validate the nanobody stabilization, the livers were isolated after imaging and analyzed by IVIS Spectrum and confocal microscopy (**Fig. 8d**). IVIS Spectrum revealed miRFP732nano signals in EGFP-expressing liver tissue. At the cellular level, confocal imaging showed extensive co-localization between the EGFP and miRFP732nano signals, supporting target-dependent stabilization of the NIR-Fb(732)_GFP_ nanobody specifically in EGFP-positive cells. This demonstrated that miRFP732nano-based NIR-Fbs retain its target-dependent stabilization *in vivo,* enabling spatially confined detection of target-expressing tissues, while SWIR imaging provided reduced autofluorescence, allowing deeper visualization of molecular targets.

## Discussion

Far-red-absorbing CBCRs^13^ combine naturally tetrapyrrole photochemistry with a compact, single-domain architecture, making them attractive templates for engineering red-shifted NIR probes. Here, we evolved the BV-binding JSC1_58120g3 domain into three small monomeric miRFP729nano, miRFP732nano and miRFP735nano. Their excitation/emission maxima extend the spectral range of NIR FPs beyond the previous limit of 702/720 nm, respectively. Together with earlier miRFPnano proteins^9–12^, these variants establish CBCR GAF domains as a diverse source of small, fluorescent scaffolds spanning a broad portion of the NIR spectrum (670-735 nm).

Engineering JSC1_58120g3 required balancing spectral red shift, fluorescence brightness and structural compactness. Elimination of photoswitching, repositioning of the BV-binding cysteine and shortening of the termini each affected fluorescence efficiency or spectral position, necessitating multiple rounds of rational design and molecular evolution. The frequent selection of mutations that increased brightness while producing a blue-shift further indicates that fluorescence efficiency and spectral position are strongly coupled in this scaffold. The three final variants therefore represent different compromises: miRFP729nano is the brightest in mammalian cells, miRFP735nano is the most red-shifted and photostable but substantially dimmer, and miRFP732nano provides an intermediate balance that made it suitable for broader characterization.

Interesting finding is that the utility of a NIR FP cannot be predicted solely from its molecular or cellular brightness. miRFP732nano is dimmer than miRFP718nano under NIR-I detection, but its larger fraction of emission beyond 1,000 nm provides a substantial advantage in the SWIR range. At equal protein concentrations, miRFP732nano produced more SWIR signal than miRFP718nano despite its lower brightness. Thus, performance of NIR FPs is detection-range dependent: miRFP732nano is better suited to off-peak SWIR imaging. Excitation closer to the miRFP732nano absorption peak may further improve its SWIR performance, as the 680 nm excitation used here favored miRFP718nano.

The compact architecture and closely positioned termini of miRFP732nano also support applications beyond whole-cell labeling. It tolerated terminal and internal fusions and could be inserted into a nanobody scaffold to generate the target-stabilized NIR-Fb. These findings extend the NIR-Fb platform into a more red-shifted spectral range and suggest that miRFP732nano may serve as a modular component of other conditionally stabilized nanobodies. Similarly, NF-κB-and AP-1-responsive constructs demonstrated that miRFP732nano can function as a transcriptional reporter for longitudinal monitoring of cellular signaling. Such reporters measure integrated pathway activity over hours rather than rapid signaling events, but their low basal signal and genetically encoded specificity make them suitable for repeated *in vivo* imaging.

Simultaneous observation of cells labeled in different colors is crucial for studying cell physiology and interactions across tissues and depths. We demonstrated multicolor, cellular-resolution 3P microscopy of miRFP732nano-and EGFP-expressing cells using single-laser excitation at 1300 nm. Using the AAV-based approach described above, we found that miRFP732nano was sufficiently bright in transduced neurons to enable imaging up to ∼950 µm and ∼300 µm from the cortical and spinal surfaces, respectively. miRFP732nano was well tolerated in target cells 1-2 months after AAV delivery, as assessed by the morphology of surrounding microglia, a sensitive readout of tissue perturbations.^41, 42^ miRFP732nano therefore provides a valuable addition to the limited list of red-shifted FPs suitable for multicolor 3P imaging *in vivo*.

At the macroscopic scale, the red shift of miRFP732nano increased its SWIR signal and enabled noninvasive imaging of skeletal muscle, inflammatory signaling and intracellular targets. SWIR detection consistently provided sharper images and lower background than NIR-I detection. The reduction in autofluorescence was particularly important in the abdomen, where gastrointestinal contents and surrounding tissues produced substantial background in the NIR-I range but little detectable signal in SWIR.

We further reduced tissue light-scattering by combining SWIR detection with 4-aminoantipyrine-based optical clearing. In scattering phantoms, SWIR detection and clearing acted complementarily, producing substantially sharper fluorescence profiles than either NIR-I imaging or SWIR imaging without clearing. The same relationship was observed *in vivo*: optical clearing increased fluorescence from miRFP732nano-expressing muscle and inflammatory reporters, whereas SWIR detection retained an intrinsic advantage in image sharpness and background suppression. Notably, SWIR imaging of the liver without clearing achieved a contrast-to-noise ratio comparable to that obtained by NIR-I imaging after clearing, while clearing recovered additional signal attenuated by the skin and abdominal wall. To our knowledge, this is the first demonstration of absorption-based *in vivo* tissue clearing paired with a genetically encoded, SWIR-compatible FP. Because the clearing approach modifies the tissue rather than the fluorescent probe, it should be applicable to other red-shifted NIR FPs.

Several limitations define directions for further studies. The emission maxima of the developed proteins are still within the NIR-I range, and SWIR imaging relies on detection of their long-wavelength emission tails. Consequently, only a fraction of emitted photons is collected beyond 1,000 nm. In addition, the progressive reduction in quantum yield and cellular brightness from miRFP729nano to miRFP735nano suggests that further red shifting is accompanied by increased nonradiative relaxation or less favorable BV–protein interactions. Structural analysis of these variants and screening of additional far-red CBCR GAF domains may help uncouple red shift from loss of brightness. The benefits of topically applied clearing agents is also limited by diffusion, with the strongest effects expected in skin and other superficial tissues. Their performance may vary among anatomical sites, and performance during repeated or chronic imaging will require further study.

In conclusion, miRFP729nano, miRFP732nano and miRFP735nano extend the compact genetically encoded NIR palette toward longer wavelengths. miRFP732nano provides a useful balance between red-shifted emission and fluorescence performance, enabling applications ranging from protein tagging and target-dependent reporters to 3P microscopy and noninvasive SWIR imaging. More broadly, this work establishes far-red CBCR GAF domains as promising templates for next-generation NIR probes and shows that combining probe engineering with modulation of the tissue optical environment can improve fluorescence imaging in living mammals.

## Methods

### Ethical statement

All live animal procedures were performed in accordance with the National Institutes of Health guidelines and approved by the Salk Institute’s and Duke University’s Institutional Animal Care and Use Committees.

### Generation of the initial template for molecular evolution

Nucleotide sequence encoding the JSC1_58120g3 was composed based on the amino acid sequence in the crystal structure (PDB: 6XHG) ^13^. Then optimization was performed with OptimumGene algorithm from GenScript for expression in human cells. The latter may include adjustments of codon usage bias and codon pair usage bias, optimization of GC content, CpG dinucleotide frequency, elimination of mRNA secondary structure and cryptic splice sites, removal of repetitive or unstable sequences. Optimized gene was synthesized by GenScript and inserted to a pBAD/His-D plasmid ^31^ (modification of pBAD/His-B vector (Invitrogen), in which a linker between N-terminal 6xHis-tag and inserted coding sequence was shortened) by KpnI/EcoRI sites, for arabinose-induced *E. coli* expression.

### General workflow of directed molecular evolution and rational design

Randomly mutated genes were introduced into *E. coli* cells, where expression was induced. High-throughput screening of mutants with enhanced fluorescence brightness was then performed using fluorescence-activated cell sorting (FACS). Selected cells were plated on agar with inductors and screening of bacterial colonies by brightness and red shift of fluorescence by macro-imaging was performed. Colonies exhibiting promising fluorescence were additionally streaked to compare their fluorescent properties side-by-side.

Mutant genes selected in *E. coli* were sequenced, and deduced protein sequences were aligned to identify positions where beneficial mutations most frequently occurred. Recurrent mutation sites were subsequently mapped onto the 3D structure of JSC1_58120g3 (PDB: 6XHG) for structural analysis. Selected clones were subcloned into a mammalian expression vector and tested in HeLa cells by flow cytometry to identify those that retain desirable properties in both bacterial and mammalian systems.

Amino acid positions identified as mutation hotspots underwent site-saturation mutagenesis using oligonucleotides with degenerate (NNS) codons (Eurofins Genomics). Top-performing mutants were pooled and used as composite PCR templates for subsequent rounds of random or site-directed mutagenesis. Random mutagenesis was complemented by steps of rational design to enable fine-tuned and targeted modification of the protein’s photophysical and biochemical properties. All steps involving protein expression were carried out at 37 °C to mimic physiological temperature of mammal organisms during the selection process, thereby promoting proper folding in mammalian cells.

### Generation of mutant libraries and expression in bacteria

Random mutagenesis was performed by error-prone PCR using the Mutazyme II polymerase mix (GeneMorph II Random Mutagenesis Kit, Agilent Technologies) according to the manufacturer’s instructions, optimized to generate up to 16 mutations per 1000 base pairs. The resulting mutated PCR products were inserted into the pBAD/His-D vector and introduced by electroporation into *E. coli* LMG194 cells harboring the pWA23h plasmid, which expresses the *hmuO* heme oxygenase from *Bradyrhizobium ORS278* under the control of a rhamnose-inducible promoter.

Mutant libraries typically contained 10^6^–10^8^ insert-bearing *E. coli* cells immediately after transformation. Libraries yielding fewer clones were discarded. Transformed LMG194 cells were grown overnight at 37 °C in LB medium supplemented with ampicillin and kanamycin and used for subsequent protocols the following day. Large libraries (≥10⁸ clones) were preserved at –80 °C in a 1:2 mixture of 50% glycerol and fresh LB medium for repeated screening.

For expression of the mutated genes, 1 ml of the overnight culture (or 3 ml of thawed glycerol stock) was inoculated into 100 ml of fresh LB medium supplemented with ampicillin, kanamycin, and 0.02% L-rhamnose. After the culture reached an OD₆₀₀ of 0.3–0.5, protein expression was induced by adding 0.004% L-arabinose, followed by incubation for 12–16 h with continuous shaking.

### Screening mutants in E. coli cells by flow cytometry

Before sorting, bacterial cells were centrifuged form LB medium, resuspended and diluted in phosphate-buffered saline (PBS) to OD_600_ of 0.03. Fluorescence-activated cell sorting was performed using MoFlo XDP (Beckman Coulter) equipped with a solid state 721 nm laser (CNI, China). The fluorescence of bacterial cells was detected using a combination of 647 nm long-pass (Semrock) and 740 nm long-pass (Chroma) filters.

### Fluorescence macro-imaging and spectral unmixing

Cells collected by FACS in SOC medium were incubated for 1 h at 37 °C with shaking (220 rpm), then plated on LB agar supplemented with 50 µg/ml kanamycin, 100 µg/ml ampicillin, 0.02% L-rhamnose, and 0.01% L-arabinose. Plates were incubated overnight at 37 °C. In some cases, bacteria were also streaked on a nitrocellulose membrane, placed over agar with same inductors and antibiotics.

Screening of bacterial colonies and streaks on agar was performed with IVIS Spectrum imaging system (Revvity) in the two following channels: 710/15 nm excitation filter and 760/20 nm emission filter (for detection of the enhancement of fluorescent brightness) and 745/15 nm excitation filter and 800/20 nm emission filter (detection of mutants with increased red-shift of fluorescence). Selected colonies were further streaked on agar and re-tested the same way as colonies. In some cases, bacterial streaks were also imaged by Leica M205 FA fluorescence stereomicroscope equipped with Cy5.5 filter set.

For spectral unmixing, *E. coli* LMG194 host cells (Invitrogen) were co-transformed with the pWA23h vector encoding heme oxygenase (hmuO gene) from *Bradyrhizobium sp.* strain ORS278, under the control of the rhamnose promoter, and corresponding pBAD vectors expressing miRFP670nano3, miRFP704nano, miRFP718nano, or miRFP732nano. Resulted strains were streaked onto a nitrocellulose membrane placed on LB agar supplemented with inductors and antibiotics. After 24 h of incubation at 37 °C, the bacterial streaks were imaged using the IVIS Spectrum CT2 across 16 spectral channels, with excitation/emission combinations including: 1. 640/15 and 680/20 nm, 2. 640/15 and 700/20 nm, 3. 647/15 and 720/20 nm, 4. 675/15 and 720/20 nm, 5. 640/15 and 740/20 nm, 6. 675/15 and 740/20 nm, 7. 640/15 and 760/20 nm, 8. 675/15 and 760/20 nm, 9. 710/15 and 760/20 nm, 10. 640/15 and 780/20 nm, 11. 675/15 and 780/20 nm, 12. 710/15 and 780/20 nm, 13. 675/15 and 800/20 nm, 14. 710/15 and 800/20 nm, 15. 675/15 and 820/20 nm, 16. 710/15 and 820/20 nm. Spectral unmixing was performed using the built-in software functionality provided with the IVIS Spectrum.

### Protein expression and purification

LMG194 cells were transformed both with pBAD plasmid encoding the mutant of interest under the control of arabinose promoter, and the pWA23h plasmid. Typically, 500 ml of fresh LB medium supplemented with 50 µg/ml kanamycin, 100 µg/ml ampicillin, and 0.02% L-rhamnose was inoculated with 5 ml of an overnight *E. coli* culture and incubated at 37 °C upon continuous shaking (220 rpm) until the cell suspension reached an OD_600_ of 0.3-0.5. Protein expression was then induced by the addition of 0.01% L-arabinose and continued at 37 °C for 12-16 h.

After expression, bacterial cells were pelleted by centrifugation at 5,000 g for 15 min at 4°C (Sorvall RC5C Plus Refrigerated Centrifuge, Marshall Scientific) and resuspended in ice-cold PBS containing 10 mg/mL lysozyme, 1 mM PMSF, and 10 mM imidazole. The cells were then disrupted by sonification (Model 3000 Ultrasonic Homogenizer; BioLogics). The insoluble fraction of lysate was removed by centrifugation at 20,000 × g for 15 min at 4 °C, and the supernatant was loaded onto Ni-NTA columns pre-equilibrated with PBS containing 10 mM imidazole. After the lysate had completely passed through the resin, 20 column volumes of PBS with 10 mM imidazole were applied, followed by 40 column volumes of PBS without additives. Elution was performed using PBS containing 100 mM EDTA.

For removal of the 6×His tag, miRFP732nano protein bearing an upstream N-terminal TEV protease cleavage site was expressed in *E. coli* and purified using Ni-NTA agarose, then transferred into TEV protease buffer (50 mM Tris-HCl, 150 mM NaCl, 0.5 mM EDTA, 1 mM DTT, pH 8.0) using PD-10 Sephadex columns. Cleavage was performed by adding 10 units of TEV protease (NEB, Cat. #P8112S) per 15 µg of protein, and incubation was performed overnight at 4 °C. The following day, the sample was exchanged into PBS using PD-10 columns and passed through fresh Ni-NTA resin to remove non-cleaved protein, free 6×His tags, and His-tagged TEV protease. The resulting protein sample was concentrated to approximately 23 mg/ml and supplemented with 10 mM β-mercaptoethanol.

### Crosslinking with glutaraldehyde and SDS-PAGE electrophoresis

miRFP732nano with removed 6×His-tag was diluted to approximately 1 mg/ml in PBS and 0.125% of glutaraldehyde was added to protein solution and immediately mixed. After 5 min incubation at room temperature, 111 mM glycine was added to stop the reaction. Then samples were supplemented with a standard loading Laemmli buffer, heated in boiling water for 3 min and 3 µg of protein was added to wells of Mini-PROTEAN TGX precast PAGE gels (BioRad, Cat. #4561094). Electrophoresis was performed in 1× Tris/Glycine/SDS buffer (BioRad, Cat. #1610732) at 100 mV for 120 min. Gel staining was performed with GelCode Blue Safe Protein Stain (ThermoFisher Scientific, Cat. #860957) according to manufacturer recommendations.

### Native PAGE electrophoresis

miRFP732nano with removed 6×His-tag, was supplemented with 12.5% glycerol and applied to wells of Mini-PROTEAN TGX precast PAGE gels (BioRad, Cat. #4561094). 5.97 and 233 µg of protein were loaded per well of a native gel to create a low and high protein gel concentrations (approximately 46.84 µM and 0.94 mM respectively). Electrophoresis was performed in 1× Tris/Glycine/ buffer (BioRad, Cat. #1610734) at 120 mV for 280 min. Gel staining was performed with GelCode Blue Safe Protein Stain (ThermoFisher Scientific, Cat. #1860957) according to manufacturer recommendations.

### Absorption and fluorescence spectroscopy

Absorbance spectra were recorded using Hitachi U-2000 spectrophotometer. Fluorescence emission spectra were acquired on an Edinburgh FLS920 fluorescence spectrometer equipped with a 450 W Xenon arc lamp (Edinburgh Xe900 450) as the excitation light source and an extended-red sensitive PMT (Hamamatsu R2658P, spectral range: 200–1010 nm) and NIR-PMT (Hamamatsu H10330-75, spectral range: 950–1700 nm) for detection. The molar extinction coefficients were determined as the ratio of the maximum absorbance at the Q-band to that at the Soret band, assuming an extinction coefficient of 39,900 M⁻¹×cm⁻¹ at the Soret band ^43^. Fluorescence quantum yields were determined relative to that of miRFP718nano ^12^ and miRFP720 ^14^ NIR FPs with known quantum yield values.

### Construction of mammalian plasmids

For testing in HeLa cells, mutated genes (without 6xHis-tag, but prserving stop-codons) were recloned from pBAD/His-D plasmid to a pcDNA3.1/myc-HisA plasmid (Invitrogen) by KpnI/EcoRI sites.

For expression in murine body and brain cells, miRFP732nano gene was inserted into an AAV-based vector with CAG promoter ^44^ by KpnI/EcoRI sites. For labeling soma of the neurons, RiboL1 peptide was fused to C-terminus of miRFP732nano and inserted into AAV-based vector with synapsin promoter ^33^ by BamHI/EcoRI sites.

To engineer plasmids for protein tagging and labeling of intracellular structures, the miRFP718nano gene was swapped with miRFP732nano either as a C-terminal fusion (for β-actin), N-terminal fusions (for mitochondria and histone H2B), or as internal fusions (for β2-adrenergic receptor (β2AR) and G-protein α-subunit (Gαs)). To construct the NIR-Fb(732) specific for GFP, the miRFP670nano3 gene was swapped with miRFP732nano in the previously described plasmid encoding NIR-Fb_GFP_ ^10^.

To construct the miRFP732nano-based NF-κB reporter, the gene encoding miRFP718nano was replaced with miRFP732nano in the miRFP718nano-based inflammation-reporter plasmid. The AP-1 reporter was then generated by replacing the NF-κB response elements in this construct with AP-1 response elements (tgagtcagtgactcagtgagtcagtgactcagtgagtcagtgactcag sequence).

To design a pAAV-shMBVR-HO1-Fd-Fnr-CW3SL construct, the short CAG promoter from pAAV-CAG-mBaoJin-oPRE plasmid was transferred to pAAV2/9-shMBVR-HO1-Fd-Fnr-CW3SL in place of CaMKII promoter by SpeI/NheI sites. pAAV-CAG-mBaoJin-oPRE, encoding bright monomeric green FP mBaoJin derived from StayGold ^45^, was obtained from Addgene (plasmid #218684).

### Mammalian cell culture and flow cytometry

HeLa and NIH3T3 cells were obtained from the ATCC and maintained in DMEM containing 4 g/L glucose, 4 mM L-glutamine, and 110 mg/l sodium pyruvate (Corning), supplemented with 10% FBS (Corning).

Transfection was performed in 12-well plates with 1:10 mix (by mass) of pEGFP-N1 plasmid and plasmid, encoding the protein of interest, using a standard PEI protocol ^46^. Cells were used for experiments after 48 or 72 h of expression. Prior to flow cytometry analysis, cells were washed twice with PBS and detached by incubation in 20 mM EDTA in PBS for 10 minutes. Subsequently, 1% BSA was added to the PBS-EDTA solution, and the cell suspension was pipetted and passed through 40-µm mesh caps into flow cytometry tubes.

Cell fluorescence intensity was analyzed by BD LSRII flow cytometer. A 637 nm laser was used for excitation of mutant proteins and emission was collected using a 740 nm long-pass filter (Chroma). For analysis of EGFP fluorescence, a 488 nm blue laser was used for excitation and 500-575 nm filter for registration of emission. For comparison of effective brightness in cells, fluorescence intensity was normalized to the excitation efficiency of each NIR FP by the 637 nm laser, and to the portion of emission spectrum of each FP in the 740 nm emission filter.

In another set of experiments, cell fluorescence was analyzed on BD Accuri C6 flow cytometer equipped with the 488 nm, and 640 nm lasers. NIR FPs were excited with 637 nm laser and emission detected with a 670 nm LP or 675/25 nm filters. EGFP was excited with a 488 nm laser, and its fluorescence was detected with a 510/15 nm emission filter.

For each sample, 50,000–100,000 events were analyzed. EGFP-positive cells were first gated, and the NIR fluorescence intensity within this gate was then assessed. Data were analyzed using FlowJo software (version 7.6.2).

### Wide-field fluorescence microscopy

Imaging of live HeLa cells was performed by using Olympus IX81 inverted microscopes, each equipped with either a 300 W xenon arc lamp (Sutter) or a 200 W metal halide arc lamp (Lumen220PRO, Prior), a 60× 1.35NA oil immersion objective lens (UPlanSApo, Olympus), and either an Orca CCD camera (Hitachi) or an opiMOS sCMOS camera (QImaging). Chroma Cy5.5 filter sets were used to acquire NIR fluorescence (HQ682/12x, HQ725/50m, Q695LP, or HQ685/20x, HQ725/40m, Q695LP), and EGFP fluorescence (HQ480/40x, HQ535/40m, 505DCXR, or HQ485/20x, HQ525/30m, 505DCXR). During imaging, HeLa cells were incubated in a cell imaging medium (Life Technologies–Invitrogen) and maintained at 35 °C. The microscope was operated using either SlideBook v.6.0.8 software (Intelligent Imaging Innovations).

### Confocal microscopy

Confocal imaging of brain or spinal sections was performed on a Zeiss LSM 710 with ZEN Black software (v2011 SP7) and an Olympus 20 × 0.8 NA air-matched objective. Two-channel, 4–6 × 4–6 tiled z-stacks (∼20 images at a 1 μm axial step size) were acquired using 488 nm and 633 nm laser lines. Emission filters were set to 500–550 nm and 700–730 nm for the 488 nm and 633 nm lines, respectively. Each z-stack image had a resolution of 1024 × 1024 pixels.

### Testing pAAV-shMBVRA-HO1-Fd-Fnr-sCAGW3SL in cultured cells

NIH 3T3 cells were transfected with equal molar quantities of pcDNA plasmid encoding the miRFP732nano and pAAV-shMBVRA-HO1-Fd-Fnr-CW3SL, encoding anti-murine BVRA shRNA, HO1, Fd, and Fnr with mitochondrial targeting sequence. pEGFP-N1 was added to a total DNA mix in a mass ratio of 1:10. In control samples, AAV-CAG-mBaoJin-oPRE plasmid (or AAV vector) or empty pcDNA3.1(+) vector was used instead of pAAV-shMBVRA-HO1-Fd-Fnr-CW3SL, to compensate the total DNA amount in transfection.

### Testing of the NF-kB-induced miRFP732nano reporter in cultured cells

HEK293T cells were transfected with plasmids encoding the miRFP732nano-based NF-κB or AP-1 reporter of inflammation and EGFP (1:10) and stimulated with TNF-α or PMA in the indicated concentrations. Measurement of miRFP732nano fluorescence was performed by flow cytometry and fluorescence microscopy 72 h after transfection.

### AAV production and titration

Recombinant AAV vector encoding miRFP732nano or NIR-Fb(732)_GFP_ under CAG promoter was produced in HEK293T cells by triple-plasmid transfection. Briefly, HEK293T cells at approximately 70% confluence were co-transfected with pAAV2/8 (Addgene plasmid #112864), pHelper (Addgene plasmid #112867) and the transfer plasmid encoding either miRFP732nano or NIR-Fb(732)_GFP_ at a 1:1:1 molar ratio using polyethylenimine. Cells were harvested 72 h after transfection and lysed, and viral particles were purified using the AAVpro Purification Kit (Takara) according to the manufacturer’s instructions.

Before titer determination, viral preparations were treated with DNase to remove unpackaged DNA. Viral genome titers were quantified as DNase-resistant genome copies by quantitative PCR using SYBR Green chemistry on a QuantStudio Real-Time PCR System (Applied Biosystems, Thermo Fisher Scientific). Quantification was performed using primers targeting rsmiRFPnano: forward, 5′-CGCGTGACCATCTACAAGTT-3′; reverse, 5′-TCGTTCAGGTAGCAGTCCT-3′.

For the mouse brain and spine imaging, recombinant AAV9 production was performed by the Viral Vector Core at the Salk Institute. The recombinant AAV9-hSyn-miRFP732nano-RiboL1 and AAV9-shMBVR-HO1-Fd-Fnr-CW3SL viruses had titers of 7.74 x10^13^ GC/ml and 7.71 x10 ^13^ GC/ml, respectively.

### Animal subjects

For multiphoton imaging, we used female *Cx3cr1^GFP/+^*mice (Jackson Laboratories; stock 005582) aged ∼13.5 – 21 weeks (N=3) and 8.5 – 11 weeks (n=4) at the time of brain and spinal cord imaging, respectively (∼4.5 – 8.5 weeks and 4 – 6 weeks after stereotactic injection; two injections per brain or spinal hemisphere; two to four injections per animal). All mice were on a C57BL/6J background. Mice were group-housed, provided with bedding and nesting material, and maintained on a 12 h light-dark cycle in a temperature-controlled (22 ± 1 °C) and humidity-controlled (45–65%) environment.

For SWIR imaging, we used male BALB/cJ mice (Jackson Laboratory, stock 000651) aged ∼8-10 weeks. Mice were fed TD.94048 AIN-93M purified diet for 1 week before imaging to reduce gastrointestinal autofluorescence from standard chow. Mice were anesthetized using 5% isoflurane with supplemental oxygen at a flow rate of 1.5 l/min. The imaging region of each mouse was first shaved with an electric trimmer and subsequently treated with a depilatory cream (Nair) to remove any remaining fur. After treating with Nair for 5 min, the abdomen was cleaned using isopropyl alcohol pads. During imaging, mice were maintained under anesthesia using 1–3% isoflurane with supplemental oxygen and their body temperature was regulated using a heating pad.

### Virus stereotactic injection

The two AAVs were co-delivered stereotactically to the cortex or the lumbar spinal cord of 6-12.5-week-old or 4.5-5.5-week-old mice, respectively. For cortical injections, 0.25 μl of each diluted AAV was injected per site at the following coordinates: AP −1.5 to −2.5 mm, ML 1.45–1.5 mm, DV 0.7–1.5 mm. AAV dilutions were 1:12 for AAV9-hSyn-miRFP732nano-RiboL1 and 1:10 for AAV9-shMBVR-HO1-Fd-Fnr-CW3SL. For intraspinal delivery, the AAV mix (same volumes and dilutions as for the cortical injections) was injected into the L4–L5 lumbar spinal cord at ML 0.25–0.35 mm, DV 0.25–0.5 mm.

Surgical procedures followed established protocols.^42, 47^ Briefly, thin-wall glass pipettes were pulled on a Flaming/Brown micropipette puller (model P-97, Sutter Instrument Company). Pipette tips were cut at an acute angle under 10× magnification using a sterile technique. Tip diameters were typically 15–20 μm. Pipettes that did not produce sharp bevels or had larger tip diameters were discarded. Millimeter tick marks were made on each pulled needle to measure the volume of virus injected into the brain.

Mice were anesthetized with isoflurane (4-5% for induction; 1–1.5% for maintenance) and positioned in a computer-assisted stereotactic system with digital coordinate readout and atlas targeting (Leica Angle Two). Body temperature was maintained at 36–37 °C using a direct-current (DC) temperature controller, and ophthalmic ointment was applied to prevent the eyes from drying out. A small amount of depilator cream (Nair) was used to remove hair from the dorsal area of the injection site. The skin was cleaned and sterilized with two-stage scrubs of Betadine and 70% ethanol.

For cranial injections, a midline incision was made, beginning just posterior to the eyes and ending just past the lambda suture. The scalp was retracted, and the periosteum was cleaned with a scalpel and forceps to expose the desired hemisphere for calibrating the digital atlas and marking coordinates. Once the reference points (bregma and lambda) were positioned with the pipette needle and entered into the program, the target was set in the digital atlas. The injection pipette was then carefully moved to the target site using AP and ML coordinates. Next, the craniotomy site was marked, and an electrical micro-drill with a fluted bit (0.5 mm tip diameter) was used to thin a 0.5–1 mm diameter area of bone over the target injection site. Once the bone was thin enough to flex gently, a 30 G needle with an attached syringe was used to carefully cut and lift a small (0.3–0.4 mm) segment of bone.

For spinal injections, a small (∼10 mm) incision was made along the midline of the L4-L5 vertebra using surgical scissors. The fascia connecting the skin to the underlying muscle was removed with forceps. Skin was held back with retractors, creating a ∼25 × 40 mm exposed area. Using blunt dissection, the lateral edges of the spinal column were isolated from connective tissue and muscle. Tissue from the vertebra of interest (L4–L5) and from one vertebra rostral and one caudal to the site of spinal cord exposure was removed with forceps. The spine was then stabilized with Cunningham vertebral clamps, and any remaining connective tissue overlying the exposed vertebrae was removed with a fine spatula. Using a small sterile needle, a ∼0.3 mm opening was made in the tissue overlying the designated injection site between the L4 and L5 vertebrae.

For injection, a drop of AAV from each virus (2–3 μl) was carefully pipetted onto parafilm to fill the pulled injection needle to the desired volume. Once loaded, the injection needle slowly lowered into the tissue until the target depth was reached. Manual pressure was applied using a 30-ml syringe connected by shrink tubing, and 0.5 μl of the virus mixture was injected slowly over 5–10 min. After injection, the syringe’s pressure valve was locked. The position was maintained for approximately 10 min to allow the virus to spread and prevent backflow during needle retraction. Each animal received two injections (∼1 mm apart) in one hemisphere. Following the injections, head clamps or Cunningham clamps were removed, the muscles were approximated, and the skin was sutured along the incision. Mice were given subcutaneous Buprenorphine SR (0.5 mg/kg) and allowed to recover before being placed in their cage.

### Animal preparation for in vivo multiphoton imaging

Surgical procedures followed established protocols.^42, 48^ Briefly, for cortical imaging, mice were anesthetized with isoflurane (4–5% for induction; 1–1.5% for maintenance) on a custom surgical bed (Thorlabs). Body temperature was maintained at 36–37 °C using a DC temperature control system, and ophthalmic ointment was applied to prevent the eyes from drying out. Depilator cream (Nair) was used to remove hair over the surgical site on the head. The skin was thoroughly cleansed and disinfected with two-stage scrubs of Betadine and 70% ethanol. A scalp flap was removed with surgical scissors to expose the frontal, parietal, and interparietal skull segments. Scalp edges were attached to the lateral sides of the skull with a tissue-compatible adhesive (Vetbond, 3M). A custom-machined metal plate was affixed to the skull with dental cement (cat. no. H00335, Coltene Whaledent) to stabilize the head in a custom holder during imaging. An approximately 3-mm-diameter craniotomy was made over the AAV injection sites, and the dura was removed. A ∼1.5% agarose solution and a round No. 0 coverslip were applied to the exposed cortical tissue. The coverslip was affixed to the skull with dental cement to control tissue motion and enable imaging.

For spinal cord imaging, mice were implanted with a spinal plate under general anesthesia 5-7 days before laminectomy. Buprenorphine SR (0.5 mg/kg) was administered to minimize postoperative pain. On the day of imaging, the mice were re-anesthetized, and a laminectomy (typically 2 mm wide and 4 mm long) was performed at the T12-T13 vertebral level, corresponding to spinal segments L2–L5. The dura mater overlying the spinal cord was kept intact, and a custom-cut No. 0 coverslip was used to seal the laminectomy, creating an optical imaging window.

Animals were imaged immediately after optical window preparation while under general anesthesia.

### In vivo multiphoton microscopy

Imaging was performed using a Movable Objective Microscope (MOM; Sutter Instrument Company) equipped with a high-repetition-rate, dual-output-channel three-photon laser (WD-1300-pro, Class 5 Photonics; center wavelength: 1300 nm; repetition rate: 1 MHz; pulse width after the objective: ∼50 fs), two fluorescence detection channels (primary dichroic beamsplitter: FF775-DiO1 (Semrock); secondary dichroic beamsplitter: 565DCXR (Chroma); green emission filter: ET-525/70m (Chroma); NIR emission filters: FF01-732/68 or FF01-708/75, supplemented with a FF01-790/SP (Semrock); photosensor modules: H7422-40 and H7422-50 (Hamamatsu) for the green and NIR channels, respectively), and MPScope software (Kleinfeld lab, UCSD; v.1). The average laser power at the tissue surface was ∼3-4 mW, and it was adjusted with depth as needed to compensate for signal loss from scattering and absorption. At the excitation intensities and durations used in this study, no signs of phototoxicity, such as blebbing of labeled cells or a gradual increase in baseline fluorescence, were apparent in our recordings. An Olympus 25× 1.05 NA water immersion objective was used. Z-stacks comprised 300-475 images, acquired at 1-2 μm axial step size, with a two-frame average, 512 × 512-pixel resolution, 2.03 Hz frame rate, and 1.4x optical zoom (corresponding to a 408 μm field of view).

Data were processed, analyzed, and plotted in Fiji (v2.14.0/1.54f; SciJava), Igor Pro (v9.05; WaveMetrics), and Adobe Illustrator (v2026). xz projections were generated from xy fluorescence image stacks using the ‘Reslice’ and ‘3D Project’ functions in Fiji. Supplemental Videos were created in Igor Pro (v9.05; WaveMetrics).

### Brain and spine tissue fixation and slicing

After *in vivo* imaging, mice were euthanized in their home cages in accordance with American Veterinary Medical Association (AVMA) guidelines. Transcardial perfusion was performed with 10% sucrose followed by 4% paraformaldehyde (PFA). Brain or spinal tissue was carefully extracted and incubated in 4% PFA overnight at 4 °C. The tissue was then washed on a shaker with 1× PBS three times at room temperature (15 min per wash). A Leica VT1000S vibratome was used to prepare 40-μm-thick coronal sections. Sections were mounted and imaged directly for native GFP and AAV-driven miRFP732nano fluorescence; no immunostaining or signal amplification was performed.

### Tissue clearing dye preparation

Gel formulations of the tissue-clearing dye for *in vivo* applications were prepared by mixing low-melting-point agarose with tissue-clearing dye solutions. 4-aminoantipyrine powder (Sigma-Aldrich) was dissolved in 1× PBS to a concentration of 38% (w/w). The dye solution was then preheated in an 80 °C water bath and mixed with agarose to achieve a final agarose concentration of 10 mg ml⁻¹. After mixing, the solution was removed from the water bath and cooled at 4 °C. Following approximately 10 min of cooling, the mixture formed a transparent orange gel that remained stable at room temperature for at least 4 h.

### Tissue clearing dye application

Gel formulations containing 4-aminoantipyrine were mixed with pumice particles when enhanced skin penetration was required. A thin layer of the formulation was topically applied to the depilated skin using a cotton-tipped applicator and gently massaged for approximately 5 min, until the skin color changed from pink to red, indicating sufficient dye penetration. The pumice particles were then removed by wiping, and a pumice-free gel formulation was subsequently applied for an additional 10 min until maximal tissue transparency was achieved.

Following imaging, the dye was removed by rinsing the treated skin with warm water or saline until the skin returned to its normal appearance. After the skin was rinsed and dried, a thin layer of Neosporin was applied to minimize the risk of skin infection.

### SWIR fluorescence imaging

A laboratory-made SWIR imaging system used a high-sensitivity InGaAs camera that has a broad detection spectrum from 600 nm to 1.7 μm (Ninox 640 II, Raptor Photonics). The camera has a pixel size of 15 μm and a maximum frame rate of 120 Hz. The camera was equipped with a VIS-SWIR objective lens with a F-number of 1.8 (RPL-OESWIRECON14mmx1.8A, Raptor Photonics). The excitation light was provided by a 680 nm laser diode (M680L5, Thorlabs) equipped with a 680/10 nm bandpass excitation filter (FBH680-10, Thorlabs). The surface exciton light intensity was roughly 50 mW cm^−2^ for all the NIR-I and SWIR imaging in this study. The camera exposure time was 150 ms for NIR-I imaging and 400 ms for SWIR imaging, if not stated otherwise. The fluorescence emission filters were 750/40 nm (FBH750-40, Thorlabs) for NIR-I imaging and 1,050 nm long pass (FEL1050, Thorlabs) for SWIR imaging.

### Data analysis in SWIR imaging

The raw white-light and fluorescence images were acquired using Micro-Manager and saved as TIFF files. Further data processing was performed using MATLAB v.2024b. For fluorescence image processing, multiple fluorescence frames acquired under identical imaging conditions were averaged to reduce random noise. Dark images acquired using the same camera settings and exposure time were also averaged and subtracted from the averaged fluorescence image to remove the mean dark background and fixed electronic offset. The image fusion of the white-light image and the corrected fluorescence image was performed by adjusting the transparency of the fluorescence image overlaid on the white-light image. The contrast-to-noise ratio (CNR) was calculated as the difference between the mean fluorescence intensity within the region of interest (ROI) and the mean background intensity outside the ROI, divided by the standard deviation of the background signal. The ROI corresponding to the liver region was manually defined based on the white-light image, and the background was taken from regions outside the ROI. The mean fluorescence intensity was computed by averaging all pixel values within the ROI.

### Intramuscular AAV delivery and tissue-clearing fluorescence imaging

AAV8-miRFP732nano was administered by intramuscular injection into the gastrocnemius muscle of mice at a dose of 5 × 10¹⁰ GC in 25 µl. 21 days after injection, mice were imaged *in vivo* to assess miRFP732nano fluorescence in the injected muscle. The injected hindlimb was then processed for tissue-clearing imaging. Fluorescence images were acquired before tissue clearing, after tissue clearing with the skin intact and after tissue clearing following removal of the overlying skin. NIR and SWIR fluorescence images were collected under identical imaging conditions for comparison across the three imaging states.

### Hydrodynamic liver transfection and LPS stimulation

In vivo plasmid delivery was performed by hydrodynamic tail vein injection as previously described. Briefly, mice were anesthetized with 1–1.5% isoflurane and rapidly injected via the tail vein with plasmids encoding the NF-κB reporter, the AP-1 reporter, or a EGFP control construct. The NF-κB and AP-1 reporters were based on miRFP732nano. The injection volume was adjusted to 8% of body weight; for a typical 20-g mouse, 1.6 ml of plasmid solution at 20 µg ml⁻¹ was delivered within 7 s.

After injection, mice were returned to their cages and monitored during recovery. At 72 h after transfection, mice received an intraperitoneal injection of lipopolysaccharide (LPS; Sigma-Aldrich, cat. no. L2630) in PBS at 2 mg kg⁻¹ body weight in a total volume of 100 µl. Fluorescence imaging was performed before LPS administration and at 6, 12, 18 and 24 h after injection. At the 24-h time point, mice were imaged sequentially under three conditions: before tissue clearing, after tissue clearing with the abdominal wall intact and after removal of the overlying abdominal skin and muscle layers to expose the liver. NIR-I and SWIR fluorescence images were acquired under matched imaging conditions, and fluorescence intensity was quantified from regions of interest over the liver area.

### In vivo evaluation of antigen-dependent miRFP732nano–based NIR-Fb_GFP_ stabilization

To evaluate target-dependent stabilization of NIR-Fb(732)_GFP_ nanobodies *in vivo*, hepatic EGFP expression was first induced by hydrodynamic tail vein injection of a pcDNA-EGFP plasmid. 24 h later, mice received AAV8-NIR-Fb(732)_GFP_ by i.v. injection at a dose of 1 × 10¹¹ GC per mouse. *In vivo* fluorescence imaging was performed 4 days after AAV transduction.

Immediately after *in vivo* imaging, mice were euthanized and livers were collected for *ex vivo* analysis using an IVIS Spectrum imaging system. For confocal microscopy, the liver tissues were then embedded, cryosectioned and imaged with a Zeiss LSM 880 Airyscan Fast inverted confocal microscope. EGFP and NIR-Fb(732)_GFP_ fluorescence was acquired using 488 nm and 633 nm excitation channels, respectively.

### Statistics and reproducibility

The sample size for imaging experiments was not statistically pre-determined. For *in vivo* multiphoton microscopy, z-stacks with weak staining, low signal-to-noise ratio, or motion artifacts were excluded. For confocal imaging, slices with tears, folds, uneven staining, or weak staining were excluded. Corresponding experiments were not randomized, and investigators were not blinded to group allocation or outcome assessment. Live animal experiments included 3–4 biological replicates (mice) per group and 2-4 technical replicates (injection sites). For tissue slice experiments, findings were replicated across 2–3 biological replicates (mice) and 4 technical replicates (tissue slices). Sex as a biological variable was not considered, as the primary goal was to demonstrate the approach’s technical capabilities. No sample-size estimation was performed to ensure adequate power to detect a pre-specified effect size.

## Data availability

All data supporting the findings of this study are available within the paper and Supplementary Information. The plasmids constructed in this study, along with their full maps and nucleotide sequences, will be available at the Addgene repository.

## Accession codes

The nucleotide sequences of miRFP729nano, miRFP732nano, and miRFP735nano have been deposited in GenBank under accession numbers PV754070, PV754071, and PV754072, respectively.

## Code Availability

The image processing codes used in this study will be available at Duke Photoacoustic Imaging Lab’s GitLab page: https://gitlab.oit.duke.edu/pilab/miRFP732nano

## Acknowledgements

We thank the Flow Cytometry core Facility at Albert Einstein College of Medicine (P30 CA013330, S10 OD026833, and S10 OD032169) for cell sorting and the Salk Institute’s Viral Vector Core (P30 CA014195) and the Waitt Advanced Biophotonics Core (P30 CA014195, P30 AG068635, Henry L. Guenther Foundation, and Waitt Foundation) for assistance with virus production and confocal microscopy, respectively. This work was supported by grants GM122567 (V.V.V.), NS142167 and NS123719 (both to A.N.), EB028143, NS111039, HL166522, DK139109, DK052985, MH135932 and ES036951 (all to J.Y.) from the US National Institutes of Health, 220011 from the Jane and Aatos Erkko Foundation (V.V.V.), 360277 from the Research Council of Finland (V.V.V.), 359487 from the Research Council of Finland (K.O.T.), the NOMIS Foundation Neuroimmunology Initiative (A.N.), and the Edwards-Yeckel Research Foundation (A.N.).

## Author contributions

K.Yu.M. and O.S.O. developed and characterized proteins *in vitro*, in bacterial and mammalian cells. K.O.T and O.S.O developed and characterized in cells the NF-kB and AP-1 reporters. Y.X., J.L. and J.Y., characterized protein performance in SWIR, performed tissue clearing, and whole-body fluorescence imaging in mice. E.C. and A.N. performed multiphoton microscopy in the brains of mice. J.Z. installed custom lasers, set up FACS sorters, and guided sorting of libraries. G.H. provided advices and training on tissue clearing. J.Y. directed the SWIR imaging and tissue clearing experiments. V.V.V. conceived, planned, and directed the overall project. All authors contributed to designing experiments, data analysis, and manuscript writing.

