## Supplementary information for "Compact red-shifted near-infrared fluorescent proteins enable deep-tissue SWIR imaging with *in vivo* optical clearing"

```
                   1        10        20         30        40        50                      60        70             85
WT JSC1_58120g3    MEQALNRVI T KIRQVSDLESIF S TTTQEVRRLFGIERVTIYKFREDYFGDFI   T   ES      E     AGGWRKLVG  S  GWEDPYLNEHQGGRFQQNQ
Rational design 1  MEQALNRVI T KIRQVSDLESIF S TTTQEVRRLFGIERVTIYKFREDYFGDFI   T   ES      E     AGGWRKLVG  S  GWEDCYLNEHQGGRFQQNQ
R1                 MEQALNRVI S KIRQVSDLESIF S ITTQEVRRLFGIERVTIYKFREDYFGDFI  S/A  ES      E     SGGWRKLVGC/N/IGWEDCYLNEHQGGRFQQNQ
R2                 MEQALNRVI S KIRQVSDLESIF S TTTQEVRRLFGIERVTIYKFREDYFGDFIM/H/N/DESR/K/L/A/M/SAGGWRKLVG C/N GWEDCYLNEHQGGRFQQNQ
R3                 MEQALKRVIN/SKIRQVSDLESIY N TATQEVRRLFGIERVTIYKFREDYFGDFI H/M/V ES   K/T/R    AGGWRKLVG  C  SWEDCYLNKHRGGRFRQNQ
R4                 MEQALKRVIN/SKIRQVSDLESIYC/NTATQEVRRLFGIERVTIYKFREDYFGDFI   H   ES    K/S     ASGWRKLVG  C  GWEDCYLNEHRGGRFRQNQ
R5                 MEQALNRVIN/SKIRQVSDLESIF S TATQEVRRLFGIERVTIYKFREDYFGDFI   H   ES     K      VSGWRKLVG  C  GWEDCYLNEHRGGRFRQNQ
R6                 MEQALNHVIN/SKIRQMSDLESIF S TATQEVRRLFDIERVTIYKFRENYFGDII   H   ES     K      VSGWRKLVG  C  GWEDCYLKEHRGGRFRQNQ
R7                 MEQALNQVVN/SKIRQMSDLESIF S TATQEVRRLFGIERVTIYKFRENYFGDII   H   ES     K      VSGWRKLVG  C  GWEDCYLNEHQGGRFRQNQ
Rational design 2  MEQALNQVVN/SKIRQMSDLESIF S TATQEVRRLFGIERVTIYKFRENYFGDII   H   ES     K      VSGWRKLVG  C  GWEDCYLNEHQGGRFRQNQ
R8                 MEQALNRVI N KIRQVSDLESIF S TATQEVRRLFDIERVTIYKFREDYFGDMI   H   ES     K      ASGWRKLVG  C  CWHDCYLNEHQGGRFRQNQ
Rational design 3  --------- - ----MADLESIK S TLTQEIRRLLDIERVTIYKFREDYFGDFI   H   ES     K      ASGWRKLVG  S  GWEDCYLNEHQGGRFRQNR
R9                 --------- - ----MADLESIK S TLTQEIRRLLDIERVTIYKFREDYFGDFI   H   ES     K      ASGWRKLVG  I  GWEDCYLNEHQGGRFRQNR
Rational design 4  --------- - ----MADLESIK S TLTQEIRRLLDIERVTIYKFREDYFGDFI   H   ES     K      ASGWRKLVG  I  GWEDCYLNEHQGGRFRQNR
R10                --------- - ----MADLESIK S TLTQEIRRLLDIERVTIYKFREDYFGDFI   H   ES     K      ASGWRKLVG  I  GWEDCYLNEHQGGRFRQNR
R11                --------- - ----MADLESIK S TLTQEIRRLLDIERVTIYKFREDYFGDFI   H   ES     K      ASGWRKLVG  I  GWEDCYLNEHQGGRFRQNR
Rational design 5  --------- - ----MADLESIK S TLTQEIRRLLDIERVTIYKFREDYFGDFI   H   ES     K      ASGWRKLVG  I  GWEDCYLNEHQGGRFRQNR

miRFP729nano       --------- - ----MADLESIK S TLTQEIRRLLDIERVTIYKFREDYFGDFI   H   ES     K      ASGWRKLVG  I  GWEDCYLNEHQGGRFRQNR
miRFP732nano       --------- - ----MADLESIK S TLTQEIRRLLDIERVTIYKFREDYFGDFI   H   ES     K      ASGWRKLVG  I  GWEDCYLNEHQGGRFRQNR
miRFP735nano       --------- - ----MADLESIK S TLTQEIRRLLDIERVTIYKFREDYFGDFI   H   ES     K      ASGWRKLVG  I  GWEDCYLNEHQGGRFRQNR

                   86            100        110        120          130       140       150       160       170         182
WT JSC1_58120g3    PFVVDDIYLGETIWEEGKFNL Q KPKRPLTDCHI   E    ALE  S  FEVKSCAVVAIFQGQKLWGLLSAFQNSAPRHWDEAEVQLLMRVADQLGVAIQQAEYLAQ
Rational design 1  PFVVDDIYLGETIWEEGKFNL Q KPKRPLTDSFI   E    ALE  S  FEVKSCAVVAIFQGQKLWGLLSAFQNSAPRHWDEAEVQLLMRVADQLGVAIQQAEYLAQ
R1                 PYVVDDIYLGETIWEEGKFNL Q KPKRPLTNSFI   E    ALE  S  FEVKSCAVVAIFQGQKLWGLLSAFQNSAPRHWDEAEVQLLMRVADQLGVAIKHAEYLAQ
R2                 PYVVDDIYLGETIWEEGKFNL Q KPKRPLTDSFIM/Q/R/LALE  S  FEVKSCAVVAIFQGQKLWGLLSAFQNSAPRHWDEAEVQLLMRVADQLGVAIQQAEYLAQ
R3                 PYVVDDIYLGETIWEEGKYNLE/HKPKRPLTNSFI  R/L   ALEF/N/RFEVKSCAVVAIFQGQKLWGLLSAFQNSAPRHWDEAEVRLLMRVADQLGVAIQQAEYLAQ
R4                 PYVVDDIYLGETLWEEGKYNLE/HKPKRPLTDSFI   R    ALE R/N FEVKSCAVVAIFQGQKLWGLLSAFQNSAPRHWDEAEVHLLMRVADQLGVAIQQAEYLAQ
R5                 PYVVDDIYLGETIWEEGKYNL Q KPKRPLTDSFI   R    ALE  R  FEVKSCAVVAIFQGQKLWGLLSAFQHSAPRHWDEAEVQLLMRVADQLGVAIQQAEYLAQ
R6                 PYVVDDIYLGEIIWEEGEYNL Q KPKRPLTDSFI   R    ALE  R  FEMKSCAAVAIFQGQKLWGMLSAFQHSAPRHWDEAEVQLLMSVADQLGVAIQQAEYLD-
R7                 PYVVDDIYLGEIIWEEGKYNL Q KPKRPLTDSFI   R    ALE  R  FEVRSCAAVAIFQGQKLWGLLSAFQHSAPRHWDEAEVQLLMRVADQLGEAIQQAEYLD-
Rational design 2  PFVVDDIYLGEIIWEEGKYNL Q KPKRPLTDSFI   R    ALE  R  FEVRSCAAVAIFQGQKLWGLLSAFQHSAPRHWDEAEVQLLMRVADQLGEAIQQAEYLD-
R8                 PFVVDDIYLGETIWEEGKYNL Q KPKRPLTDSFI   R    ALE  R  FEVKSCAVVAIFQGQKLWGMLSAFQHSAPRHWDEAEVQLLMRVADQLGVAIQQAEYLAQ
Rational design 3  PFVVDDIYLGETIWEDGKYNL Q KPKRPLSDSFI   R    ALE  R  FEVKSCAVVAIFQGQKLWGMLSAFQHSAPRHWDEAEVQLLMRTADQLGVIIQQA-----
R9                 PFVVDDIYLGETIWEDGKYNL Q KPKCPLSDSFI   R    ALE  R  FEVKSCAAVAIFRGQKLWGMLSAFQHSAPRHWDEAEVQLLMRTADQLGVIIQQA-----
Rational design 4  PFVVDDIYLGETIWEDGKYNL Q KPKRPLSDSFI   R    ALE  R  FEVKSCAAVAIFRGQKLWGMLSAFQHSAPRHWDEAEVQLLMRTADQLGVIIQQA-----
R10                PYVVDDIYQGETIWEDGKYNL Q KPKRPLSDSFI   R    ALE  R  FEVKSCAAVAIFQGQKLWGMLSAFQHSAPRHWDEAEVQLLMRTADQLGVIIQQA-----
R11                PFVVDDIYLGETIWEDGKYNL Q KPKSPLSDSFI   R    ALE  R  FEVKSCAVVAIFRGQKLWGMLSAFQHSAPRHWDEAEVQLLMRTADQLGVIIQQA-----
Rational design 5  PFVTDDIYLGETIWEDGKYNL R KPKRPLSDSFI   R    ALE  R  FEVKSTAAVAIFQGQKLWGLLSAFQHSAPRHWDEAEVQLLMRTADQLGVIIQQA-----

miRFP729nano       PYVVDDIYQGETIWEDGKYNL Q KPKRPLSDSFI   R    ALE  R  FEVKSCAAVAIFQGQKLWGMLSAFQHSAPRHWDEAEVQLLMRTADQLGVIIQQA-----
miRFP732nano       PFVVDDIYLGETIWEDGKYNL Q KPKSPLSDSFI   R    ALE  R  FEVKSCAVVAIFRGQKLWGMLSAFQHSAPRHWDEAEVQLLMRTADQLGVIIQQA-----
miRFP735nano       PFVTDDIYLGETIWEDGKYNL R KPKRPLSDSFI   R    ALE  R  FEVKSTAAVAIFQGQKLWGLLSAFQHSAPRHWDEAEVQLLMRTADQLGVIIQQA-----
```

**Supplementary Figure 1. Alignment of protein clones with the highest brightness in live HeLa cells selected at each step of the directed molecular evolution.** Each step consisted of mutagenesis, selection of the brightest clones in bacteria, followed by selection of the brightest clone in HeLa cells. Amino acid substitutions in each selected clone relative to WT JSC1_58120g3 GAF domain are highlighted yellow. Eleven random mutagenesis rounds of the molecular evolution are indicated as **R_number**. Five ***Rational design*** rounds of the molecular evolution are the following. **1)** Repositioning the covalent bond between the protein and biliverdin IXα chromophore to eliminate photoswitching and preserve the large NIR shift of fluorescence maximum: P71C, C116S, and H117F. **2)** Substituting Y87 with F to eliminate a blue shift in fluorescence maximum. **3)** Removing the N-terminal α-helix (residues 1–16) and five C-terminal residues (178–182) to bring the N- and C-termini close. Additional mutations were introduced to stabilize the protein structure after termini truncations: F22K, A25L, V29I, G34L, Q85R, E101D, L113S, V166T, and A173I. **4)** Replacing C111 with R to prevent the formation of cysteine dimers. **5)** Introducing mutations V89T, Q107R, S111R, C129T, and M143L to achieve the large NIR shift of fluorescence maximum. Three final selected protein variants, termed miRFP729nano, miRFP732nano, and miRFP735nano, are highlighted blue.

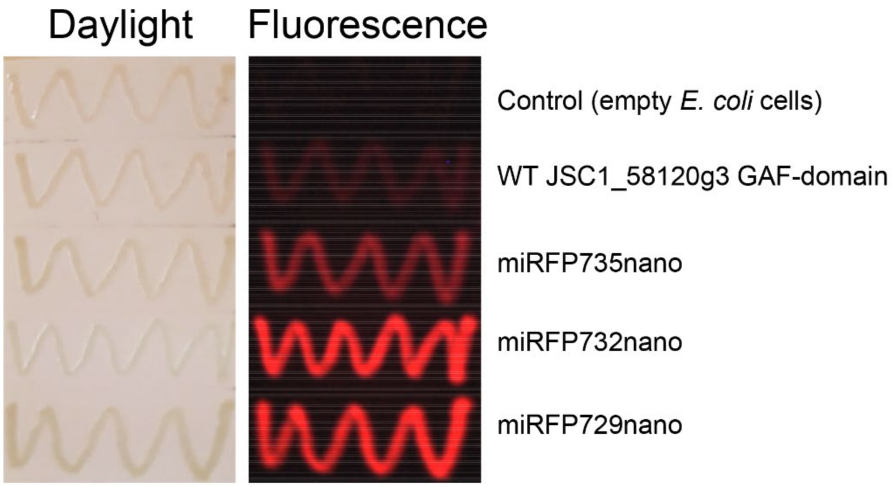


**Supplementary Figure 2. Fluorescence brightness of the parental CBCR's JSC1_58120g3 GAF domain and engineered miRFP729nano, miRFP732nano, and miRFP735nano proteins in bacteria.** Bacterial streaks co-expressing proteins and *Bradyrhizobium* hmuO heme oxygenase were photographed in day light (left) or using Leica M205 FA fluorescence stereomicroscope equipped with custom Cy5.5 (650/45 nm excitation and 710/50 nm emission) filter set (right).

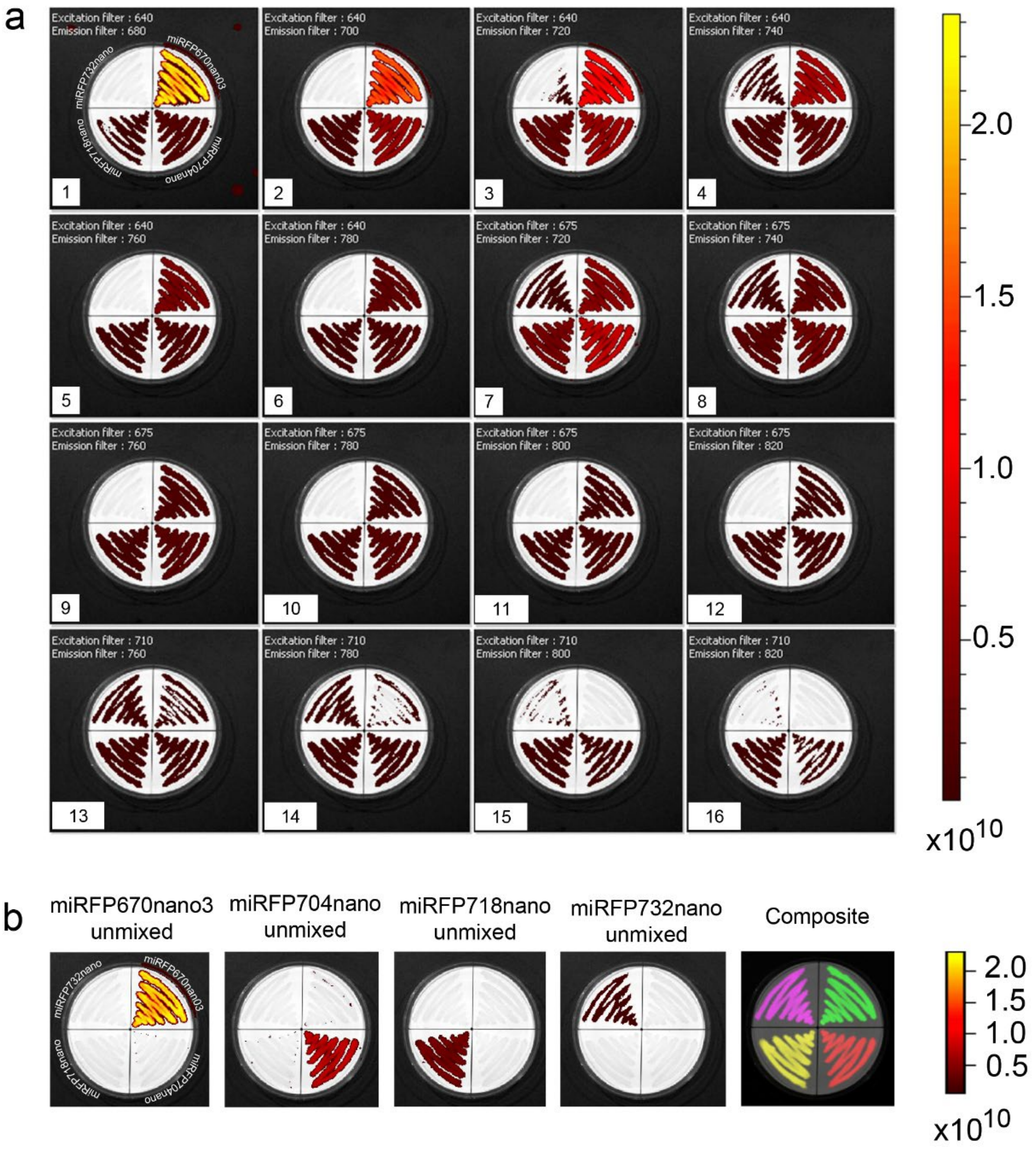


**Supplementary Figure 3. Four-color imaging of the CBCR-derived miRFPnano series of small near-infrared fluorescent proteins, including miRFP732nano. (a)** Spectral images of four bacterial streaks co-expressing different miRFPnano proteins with *Bradyrhizobium* hmuO heme oxygenase, captured in 16 spectral channels using the IVIS Spectrum imaging system. The fluorescent proteins are arranged as follows, starting from the upper right and moving clockwise: miRFP670nano3, miRFP704nano, miRFP718nano, and miRFP732nano. **(b)** Spectrally unmixed signals and composite pseudocolored images are shown (miRFP670nano3 as green, miRFP704nano as red, miRFP718nano as yellow, and miRFP732nano as magenta). Spectra from individual bacterial streaks were used as reference points for the unmixing process. The color bar represents the fluorescent radiant efficiency, defined as ([photons/s/cm²/steradian]/[µW/cm²]).

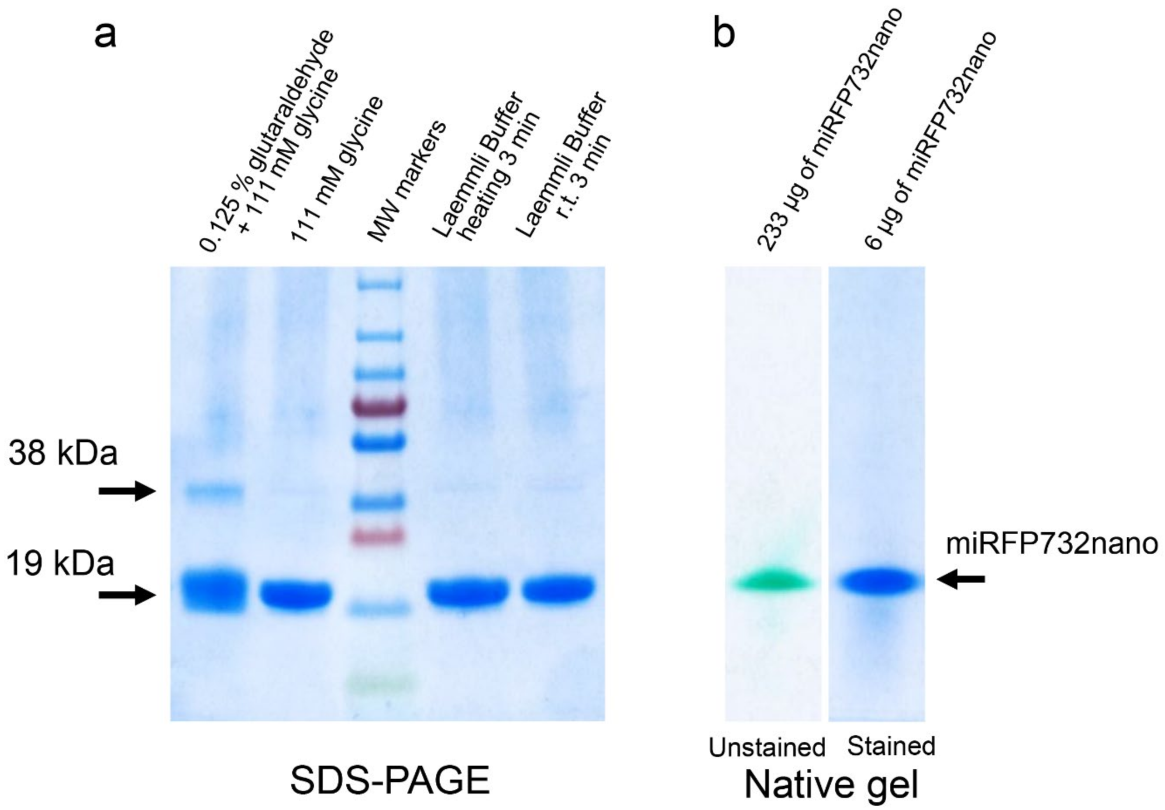


**Supplementary Figure 4. PAGE analysis of purified miRFP732nano protein lacking N-terminal 6×His-tag. (a)** SDS-PAGE gel. The 3 µg of the protein was added to each gel well. Left to right: miRFP732nano crosslinked with 0.125% glutaraldehyde, quenched by 111 mM of glycine at room temperature (r.t.); non-crosslinked miRFP732nano in the presence of glutaraldehyde quencher 111 mM of glycine at r.t. (crosslinking control); MW markers; miRFP732nano, heated at 98°C after addition of loading Laemmli buffer; and miRFP732nano, non-heated, after addition of Laemmli buffer. Crosslinking was performed for 5 min at r.t. The composition of 4×Laemmli buffer was 200 mM Tris-Cl pH 6.8, 8% SDS, 20% β-mercaptoethanol, 0.4% bromophenol blue, and 40% glycerol. The addition of 0.125% glutaraldehyde at r.t. results in the appearance of crosslinked dimer (~5%). **(b)** Native PAGE gel. Left: unstained miRFP732nano photographed in day light reveals green hue of the protein band, reflecting its effective reflection of the visible light. Right: miRFP732nano stained by Coomassie blue as in (a) but without SDS and β-mercaptoethanol added. miRFP732nano migrates as a single band even when loaded at a concentration of 0.94 mM (233 µg).

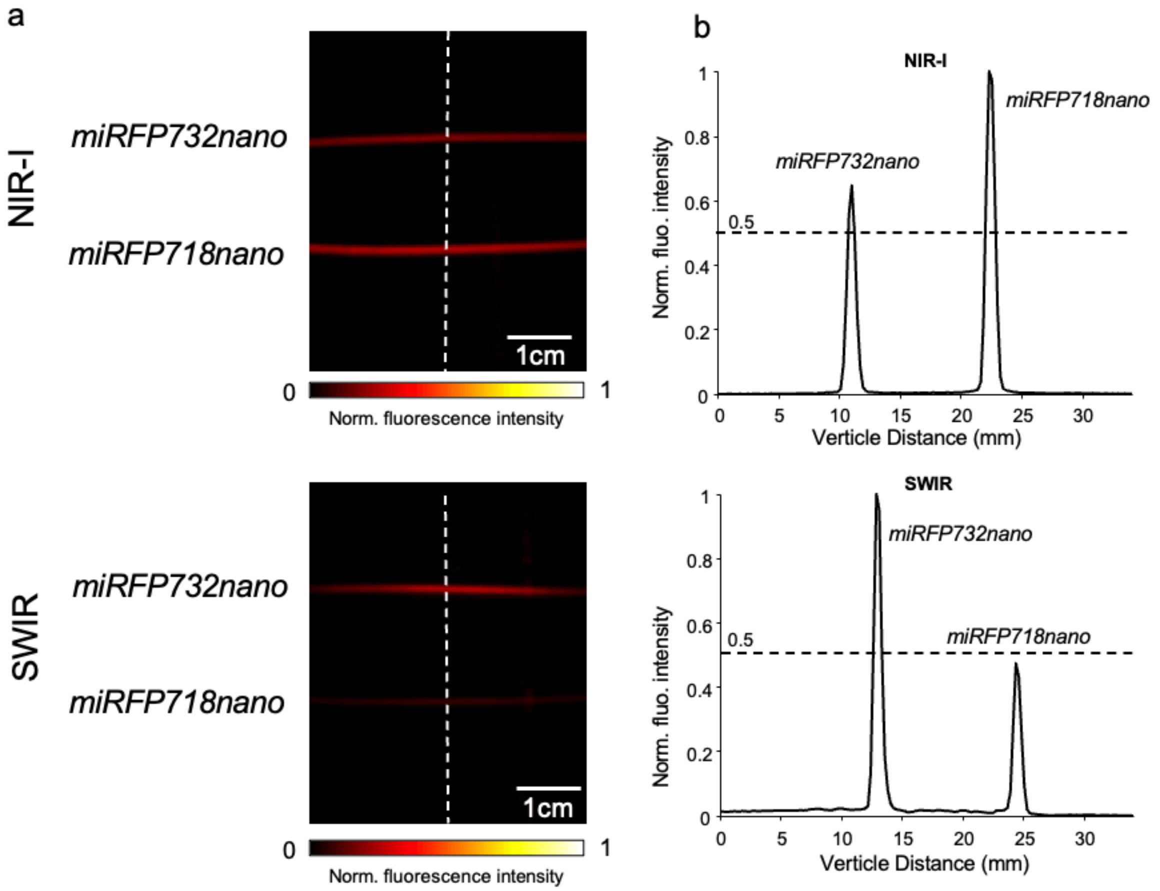


**Supplementary Figure 5**. **Comparison of miRFP732nano and miRFP718nano under NIR-I and SWIR imaging in a silicone tube phantom. (a)** NIR-I (top) and SWIR (bottom) fluorescence images of two silicone tubes (0.31 mm inner diameter) filled with 5 mg/mL purified miRFP732nano or miRFP718nano protein. Scale bars, 1 cm. **(b)** Normalized fluorescence intensity profiles along the white dashed lines in (a), under NIR-I (top) and SWIR (bottom) imaging. Dashed horizontal lines indicate the half-maximum (0.5) level.

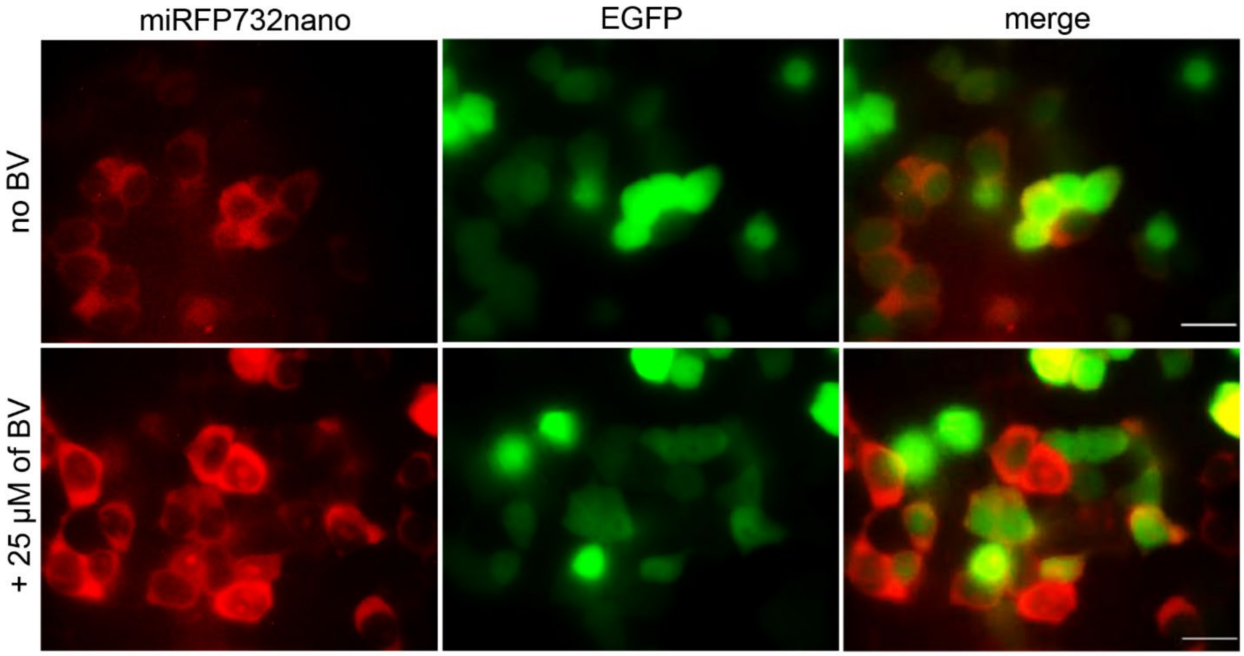


**Supplementary Figure 6. Expression of miRFP732nano fused with RiboL1 peptide encoded in an AAV plasmid under hSyn promoter.** miRFP732nano-RiboL1 fusion was inserted in a pAAV backbone bearing the WPRE, hGH poly(A) signal and two AAV2 ITR elements. Fluorescence images of live HEK293T cells co-transfected with pAAV9-hSyn-miRFP732nano-RiboL1 and pcDNA-EGFP plasmids without and with supply of exogenous BV. miRFP732nano was imaged in the Cy5.5 channel and EGFP in the FITC channel. Scale bars, 20 µm.

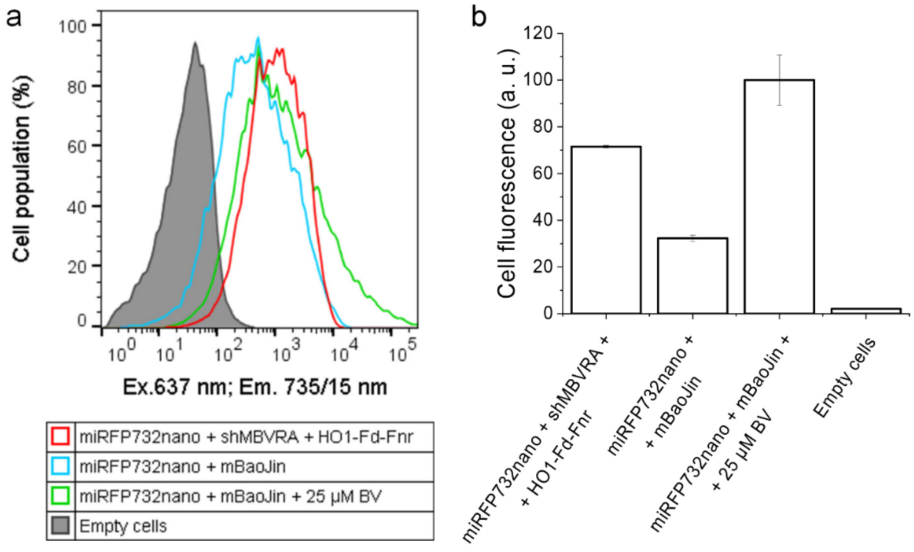


**Supplementary Figure 7. Co-transfection of NIH3T3 murine cells with pcDNA-miRFP732nano and pAAV9-shMBVRA-HO1-Fd-Fnr-*C*W3LS plasmids. (a)** Flow cytometry comparison of miRFP732nano fluorescence with and without co-expression of heme oxygenase 1 (HO1), ferredoxin (Fn), ferredoxin-NADP$^+$ reductase (Fnr), and shRNA targeting murine biliverdin reductase A (BLVRA). Cells were excited by 637 nm laser, and the fluorescence was detected at 735/15 nm filter. (b) Quantification of the data in (a). The pAAV9-mBaoJin-CW3LS plasmid, encoding monomeric version of StayGold green fluorescent protein called mBaoJin, was used to equalize the total DNA amount in the co-transfections without pAAV9-shMBVRA-HO1-Fd-Fnr-CW3LS. The mBaoJin and shMBVRA-HO1-Fd-Fnr AAV plasmids have the same short CAG promoter.

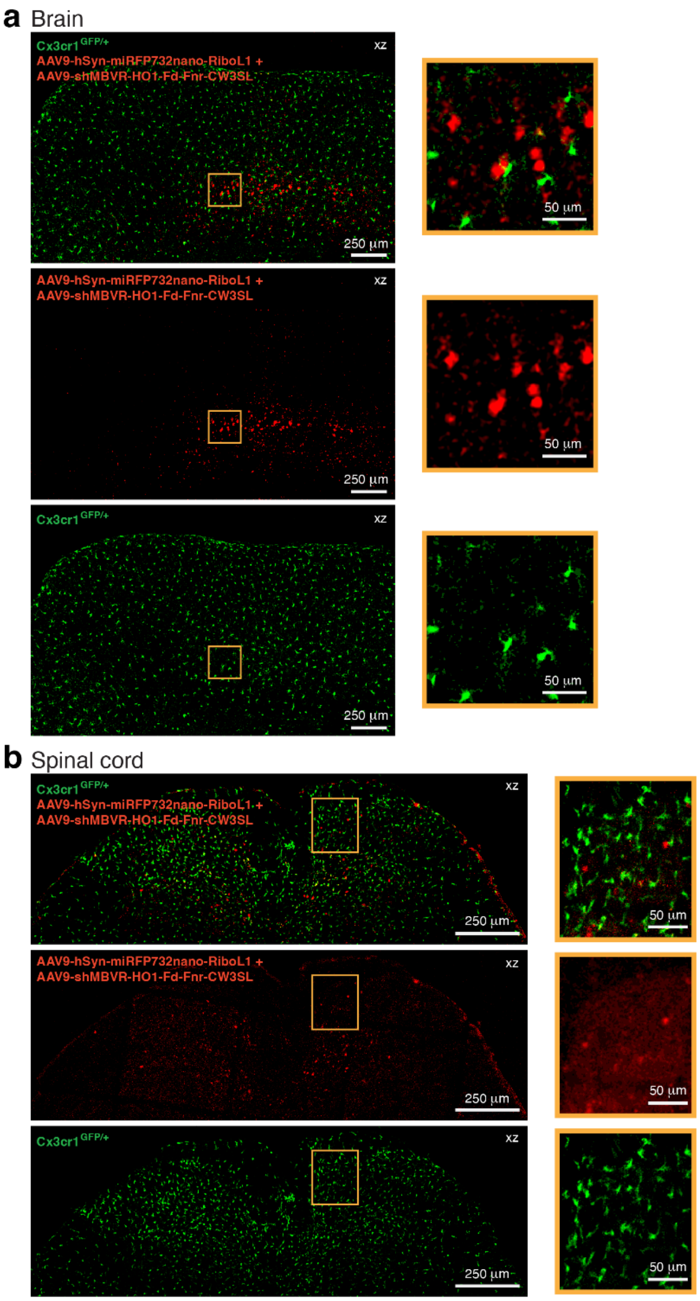


**Supplementary Figure 8. Histological analysis of miRFP732nano expression in the brain and spinal cord of *Cx3cr1*$^{GFP/+}$ mice after *in vivo* three-photon imaging.** (**a**) *Left*, confocal fluorescence images showing miRFP732nano-expressing neuronal somata (red) and GFP-expressing microglia (green) in the cortex of a *Cx3cr1*$^{GFP/+}$ mouse 4.5 weeks after stereotactic co-injection of AAV9-hSyn-miRFP732nano-RiboL1 and AAV9-shMBVR-HO1-Fd-Fnr-CW3SL viruses into deep cortical layers (*top*: overlay; *center and bottom*: color-separated images). *Right*, zoom-ins of the indicated regions. Scale bars, 250 µm (*left*) and 50 µm (*right*). (**b**) Same as in panel (a), but in a *Cx3cr1*$^{GFP/+}$ mouse 4 weeks after stereotactic co-injection of AAV9-hSyn-miRFP732nano-RiboL1 and AAV9-shMBVR-HO1-Fd-Fnr-CW3SL viruses into the lumbar spinal cord. Scale bars, 250 µm (*left*) and 50 µm (*right*). Representative images from three brain and two spinal cord histology experiments are shown.

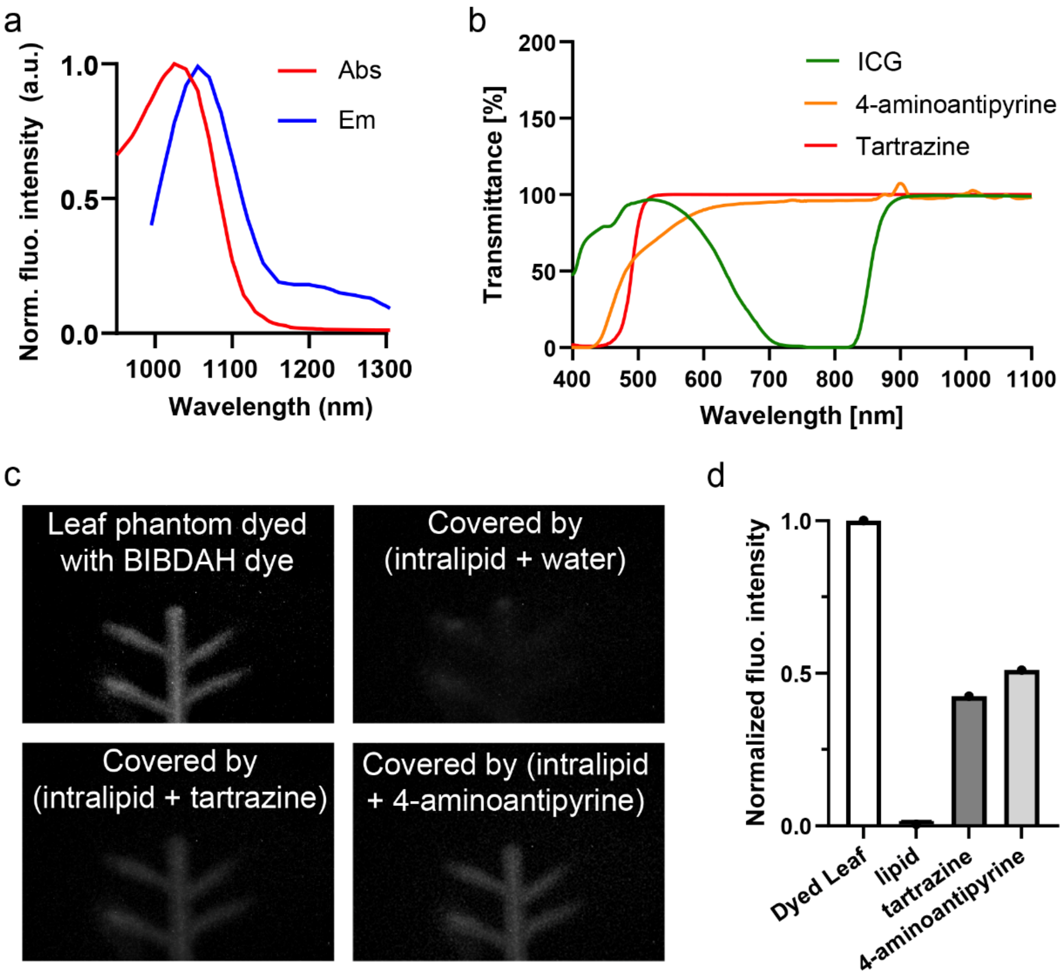


**Supplementary Figure 9 Achieving optical transparency by dye molecules in the SWIR window. (a)** Fluorescence emission of BIBDAH dye in phosphate-buffered saline. **(b)** Measured transmittance spectra of three candidate clearing dyes: ICG, tartrazine, and 4-aminoantipyrine. ICG exhibits strong absorption in the NIR-I region, leading to reduced transmission, whereas both tartrazine and 4-aminoantipyrine maintain high transmittance across the NIR-I and SWIR windows. **(c)** Comparison of clearing performance using a leaf phantom stained with BIBDAH dye. The leaf pattern is largely obscured when covered by intralipid alone and becomes visible when either tartrazine or 4-aminoantipyrine is added. However, tartrazine shows reduced clearing performance in practice due to its tendency to precipitate, which limits its effectiveness. **(d)** Quantification of normalized fluorescence intensity under different conditions, confirming that 4-aminoantipyrine achieves higher signal recovery compared to tartrazine, while intralipid alone results in minimal detectable signal.

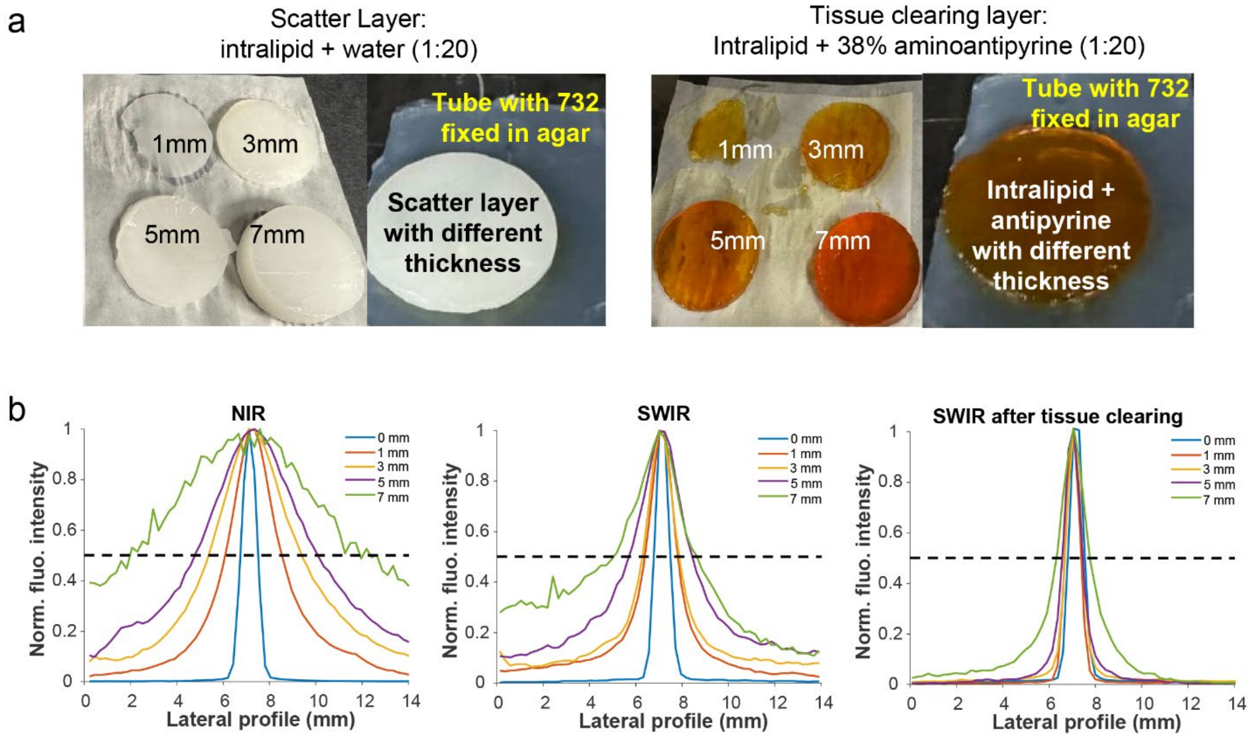


**Supplementary Figure 10. NIR-I and SWIR imaging of miRFP732nano protein in a silicone tube phantom. (a)** Photographs of the scattering layers (intralipid/water, 1:20) and tissue-clearing layers (intralipid/38% 4-aminoantipyrine, 1:20) prepared at different thicknesses (1–7 mm), used to cover a silicone tube (0.31 mm inner diameter) filled with 5 mg/mL miRFP732nano protein embedded in agar. **(b)** Normalized fluorescence intensity profiles across the tube (lateral profiles) at increasing layer thicknesses (0–7 mm), measured under NIR-I imaging (*left*), SWIR imaging (*middle*), and SWIR imaging with tissue clearing (*right*). Dashed lines indicate the half-maximum level used for FWHM determination.

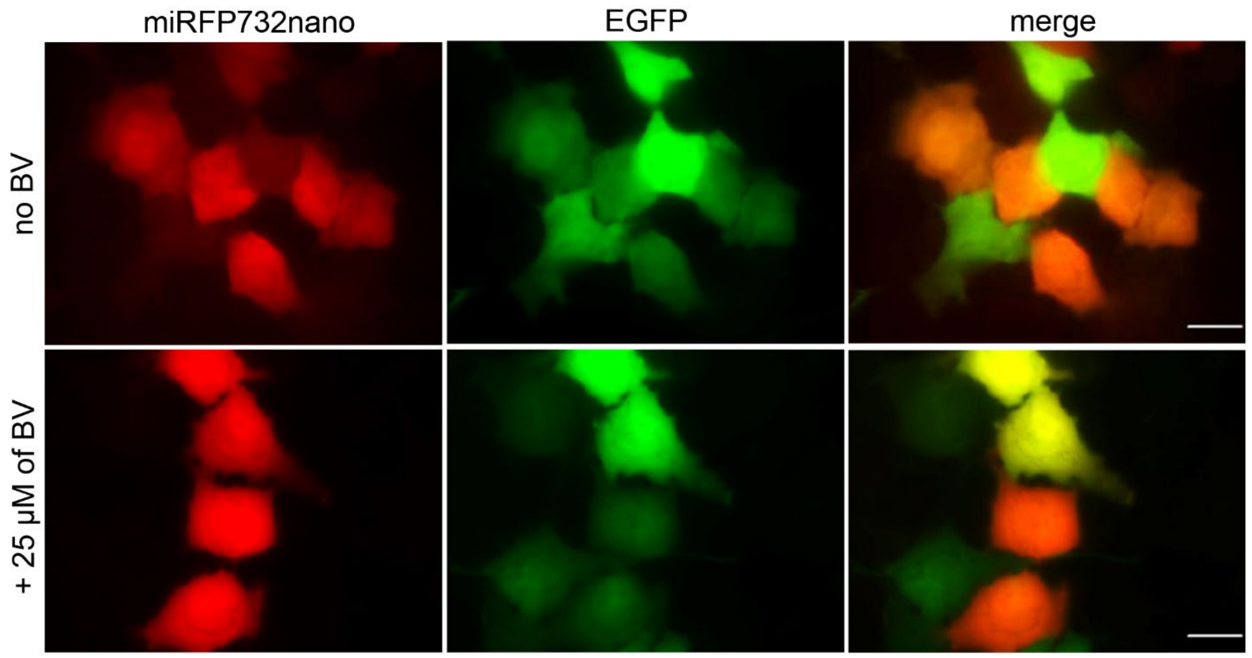


**Supplementary Figure 11. Expression of miRFP732nano encoded in an AAV plasmid under ubiquitously activated CAG promoter.** miRFP732nano was inserted in pAAV backbone bearing the WPRE, hGH poly(A) signal and two AAV2 ITR elements. Fluorescence images of live HeLa cells co-transfected with pAAV9-CAG-miRFP732nano and pcDNA-EGFP plasmids without and with supply of exogenous BV. miRFP732nano was imaged in the Cy5.5 channel and EGFP in the FITC channel. Scale bars, 20 µm.

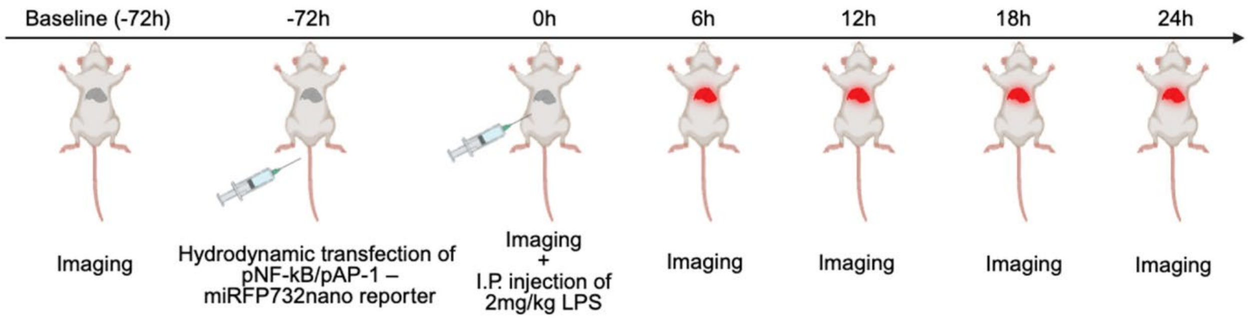


**Supplementary Figure 12. LPS-induced liver inflammation mouse model.** Schematic of the lipopolysaccharide (LPS)-induced inflammation mouse model and imaging timeline. Baseline imaging was performed 72 h prior to the LPS treatment, followed by serial imaging from 0 to 24 h after LPS treatment.

**Supplementary Table 1.**

Properties of single-domain CBCR-derived miRFPnano proteins and selected two-domain NIR FPs with emission above 700 nm.

| NIR FP | Ex, nm | Em, nm | Relative emission above 1000 nm[a] | Extinction coefficient, $M^{-1}cm^{-1}$ | Quantum yield (QY), % | Molecular brightness *vs.* miRFP670nano, % | p*K*a | Photostability in HeLa cells, $t_{1/2}$, s | Brightness in mammalian cells *vs.* miRFP670nano, % [b] | Ref. |
|---|---|---|---|---|---|---|---|---|---|---|
| miRFP670nano | 645 | 670 | 0.3 | 95,000 | 10.8 | 100 | 3.7 | 545 | 100 | 1 |
| miRFP704nano | 680 | 704 | n.d. | 93,000 | 9.8 | 90 | 4.1 | 1260 | 134 | 2 |
| miRFP718nano | 690 | 718 | 1.0 | 79,000 | 5.6 | 43 | 3.8 | 1254 | 55 | 3 |
| **miRFP729nano** | 714 | 729 | 1.9 | 69,300 | 1.9 | 13 | 4.2 | 1445 | 19 | this work |
| **miRFP732nano** | 716 | 732 | 3.5 | 64,400 | 1.6 | 10 | 4.2 | 1585 | 14 | |
| **miRFP735nano** | 719 | 735 | 3.2 | 59,200 | 1.0 | 6 | 4.1 | 1775 | 4 | |
| miRFP703 | 674 | 703 | 0.53 | 90,900 | 8.6 | 76 | 4.5 | 394 | 61 | 4 |
| miRFP709 | 683 | 709 | 0.65 | 78,400 | 5.4 | 41 | 4.5 | 192 | 42 | |
| SNIFP | 697 | 720 | n.d. | 75,000 [c] | 2.2 | 16 | 4.5 | n.d. | n.d. | 5 |
| mIFP | 683 | 704 | n.d. | 82,000 | 8.4 | 67 | 4.5 | 54 | 26 | 6, 7 |
| BDFP1.5 | 688 | 711 | n.d. | 74,000 | 5.0 | 36 | 2.0 | 1310 [c] | 0.5 [d] | 8 |

[a] Determined as integrated emission spectrum above 1000 nm normalized to the total integrated emission spectrum.
[b] Unless otherwise stated, it is determined as effective NIR fluorescence in live HeLa cells 72 h after transfection with no supply of exogenous biliverdin chromophore and after normalization to the fluorescence of co-transfected EGFP.
[c] Estimated from SNIFP absorption spectrum in Supplementary Figure 2a in the original paper[5].
[d] Based on the comparison with smURFP in HEK293 cells in[8].
n.d., not determined..
